# The Unique Role of Intracellular Perinuclear β-Adrenergic Receptors in defining Signaling Compartmentation and Pathological Cardiac Remodeling

**DOI:** 10.1101/2024.10.02.616267

**Authors:** Moriah Gildart Turcotte, Anne-Maj Samuelsson, Sofia M. Possidento, Jinliang Li, Zhuyun Qin, Michael S. Kapiloff, Kimberly L. Dodge-Kafka

## Abstract

The β-adrenergic receptor is a prototypical G-protein coupled receptor that initiates signaling from the plasma membrane. However, active receptors have been detected within intracellular compartments. The functional significance of these intracellular receptors remains unclear, including whether they regulate distinct cellular processes or function independently of plasma membrane receptors. We show using live cell imaging of primary cardiomyocytes that β-adrenergic receptors localized to Golgi apparatus opposing the outer nuclear membrane are sufficient and necessary for the stimulation of cAMP and calcium signaling within a nanometer scale compartment independent of receptors at other sites. Using compartment-specific activators and inhibitors, we show Golgi β-adrenergic receptors associated with the scaffold protein AKAP6β and the outer nuclear membrane protein nesprin-1α are responsible for pathological gene transcription and the induction of cardiomyocyte hypertrophy. The functional significance of Golgi-localized receptors is demonstrated in mice models of cardiomyopathy, providing proof-of-concept for a compartment-specific therapeutic intervention in Dilated Cardiomyopathy.

## Introduction

G-protein coupled receptors (GPCR) are critical regulators of cellular function and constitute the most common target for pharmacological invention in disease.^1^ Agonist binding to G_αs_-coupled receptors induces the production of the second messenger cAMP by transmembrane adenylyl cyclase. Foundational studies in the 1970s regarding the action of catecholamines and prostaglandins on the heart showed that activation of different GPCRs could induce distinct cAMP responses in the same cell, despite acting through a common soluble, apparently freely diffusible second messenger.^2^ It is now appreciated that cAMP signaling can be restricted to small distances (<60 nm) from the initiating GPCR.^3^ Accordingly, cAMP signaling is highly compartmentalized by the differential intracellular localization of adenylyl cyclases, phosphodiesterases, and cAMP effector proteins, often in association with multimolecular signaling complexes or “signalosomes” organized by A-kinase anchoring proteins (AKAPs).^2–4^ GPCRs are classically described as plasma membrane proteins, while GPCRs detected within the cell have been mainly considered in terms of receptor recycling or down-regulation. However, consistent with the presence of cAMP compartments at a distance from the plasma membrane, GPCRs present on intracellular membranes associated with endosomes, Golgi apparatus, sarcoplasmic reticulum, and the nuclear envelope have been suggested to directly activate signaling at sites deep within the cell.^5–9^ Whether these intracellular pools of GPCRs independently regulate specific cellular processes remains unclear, although their proximity to downstream target organelles suggests the potential for enhanced signaling specificity. In addition, while GCPRs are subject to AKAP-dependent regulation at the plasma membrane,^2^ it is not known whether GCPRs at intracellular sites are associated with AKAP signalosomes, participating directly in compartmentalized cAMP signaling. Here, we define a perinuclear GPCR signaling compartment that independently regulates the phenotype of the cardiac myocyte. We demonstrate the functional relevance of a discrete pool of intracellular GPCRs to cellular stress responses through the local regulation of a specific AKAP signalosome and its associated cAMP and Ca^2+^ signaling compartment.

The β-adrenergic receptor (βAR) regulates physiological cardiac function through the control of heart rate (chronotropy), strength of contraction (inotropy), and rate of relaxation (lusitropy). In addition, chronically elevated βAR activity in disease due to abnormally high levels of the endogenous catecholamines epinephrine and norepinephrine results in the induction of pathological cardiac remodeling.^10^ At the cellular level, this stress response includes altered myocyte contractility and metabolism, myocyte hypertrophy, increased myocyte death, and myocardial fibrosis, that together contribute to the development of end-stage heart failure.^11^ Inhibition of cardiac remodeling by β-blockade is standard of care for the prevention and treatment of heart failure, although treatment can have significant side effects.^12^ Despite guideline directed medical therapy, mortality from heart failure remains high, suggesting further investigations into the molecular mechanism and target specificity for βAR signaling is needed. cAMP and its effector protein kinase A (PKA) mediate many of the downstream effectors of βAR signaling in the myocyte, and AKAP scaffolds are known to play an important role in both cardiac physiology and pathogenesis.^4^ In cardiac myocytes the 230-kDa scaffold protein AKAP6*β* (also known as mAKAPβ) is localized to the nuclear envelope by direct binding to the outer nuclear membrane (ONM) Klarsicht, ANC-1, Syne Homology (KASH) domain protein nesprin-1*α*.^13–15^ AKAP6*β* organizes signalosomes that integrate cAMP, Ca^2+^, phosphatidylinositol 4-phosphate, mitogen-activated protein kinase, and hypoxic signaling regulating class IIa histone deacetylases (HDACs) and the transcription factors nuclear factor of activated T-cells (NFAT), serum response factor (SRF), myocyte enhancer factor 2 (MEF2), and hypoxia inducible factor 1α (HIF-1α), which in turn regulate myocyte gene expression.^16,17^ Accordingly, AKAP6β is required for the hypertrophy of primary ventricular myocytes *in vitro* and for the induction of pathological cardiac remodeling and heart failure by isoproterenol infusion, chronic pressure overload, and myocardial infarction *in vivo.*^18–23^

AKAP6 was one of the first AKAPs shown to orchestrate an independent cAMP microdomain. The scaffold binds PKA, the phosphodiesterase PDE4D3, adenylyl cyclase (AC) type 5, as well as a second cAMP effector Epac1 (Rap guanine nucleotide exchange factor 3), resulting in the local production, utilization and hydrolysis of cAMP. ^19,21,24–27^ Notably, cAMP activation of PKA bound directly to AKAP6β is required for the compartment-specific activation of the calcineurin-NFAT pathway and myocyte hypertrophy.^21,27^ Here we demonstrate that AKAP6β signalosomes are activated by βARs stimulated within the myocyte by catecholamines. Evidence provided by live cell imaging and phenotyping of primary cardiac myocytes *in vitro, shows* that AKAP6β signalosomes are regulated by βARs on Golgi membranes facing the ONM independently of other βARs at other sites in the cell, thereby defining AKAP6β-mediated cAMP signaling to a nanometer scale compartment. Importantly, while direct stimulation of βARs localized to other sites in the cell increased cAMP in the cytosol, only Golgi-localized βARs induced pathological gene transcription. These results demonstrate the functional relevance of GPCRs within a specific intracellular compartment and, moreover, suggest a new paradigm for therapeutic GPCR targeting. Targeting of GPCRs associated with individual signalosomes should affect cellular responses with fewer on-target adverse effects than that currently associated with traditional GPCR antagonists. Demonstration of the functional relevance of βARs within the Golgi-ONM compartment is corroborated by *in vivo* gain and loss of function of these receptors in mouse models of cardiomyopathy, providing proof-of-concept that targeting of AKAP6β-associated βARs might be beneficial in disease.

## Results

### An intracellular pool of β-adrenergic receptors activates AKAP6β-bound PKA

Compartmentation of cAMP-PKA signaling is conferred in part by association with AKAPs restricted to specific sites within the cell. AKAP6 is localized to the ONM by nesprin-1α.^13–15^ In past live cell imaging studies of AKAP6-bound PKA activity, we have utilized the ratiometric Fluorescence Resonance Energy Transfer (FRET) biosensor AKAR4-nesprin, herein termed “ONM-AKAR4” (Figure 1A,B), which is comprised of the PKA activity reporter AKAR4 in fusion to nesprin-1α and has a similar dynamic range as the parental AKAR4 sensor.^28^ Importantly, in neonatal and adult rat ventricular myocytes and hippocampal neurons, perinuclear PKA activity detected by ONM-AKAR4 has been shown to be dependent upon AKAP6 expression, while the diffusely localized cytosolic parent AKAR4 sensor is AKAP6 independent.^27,28^ The nuclear envelope is distant from the plasma membrane where βARs are canonically activated. Here we employed the AKAR4 sensors to test the hypothesis that an intracellular pool of βARs selectively activates AKAP6β-associated perinuclear cAMP-PKA signaling in the cardiac myocyte.

**Figure 1:**
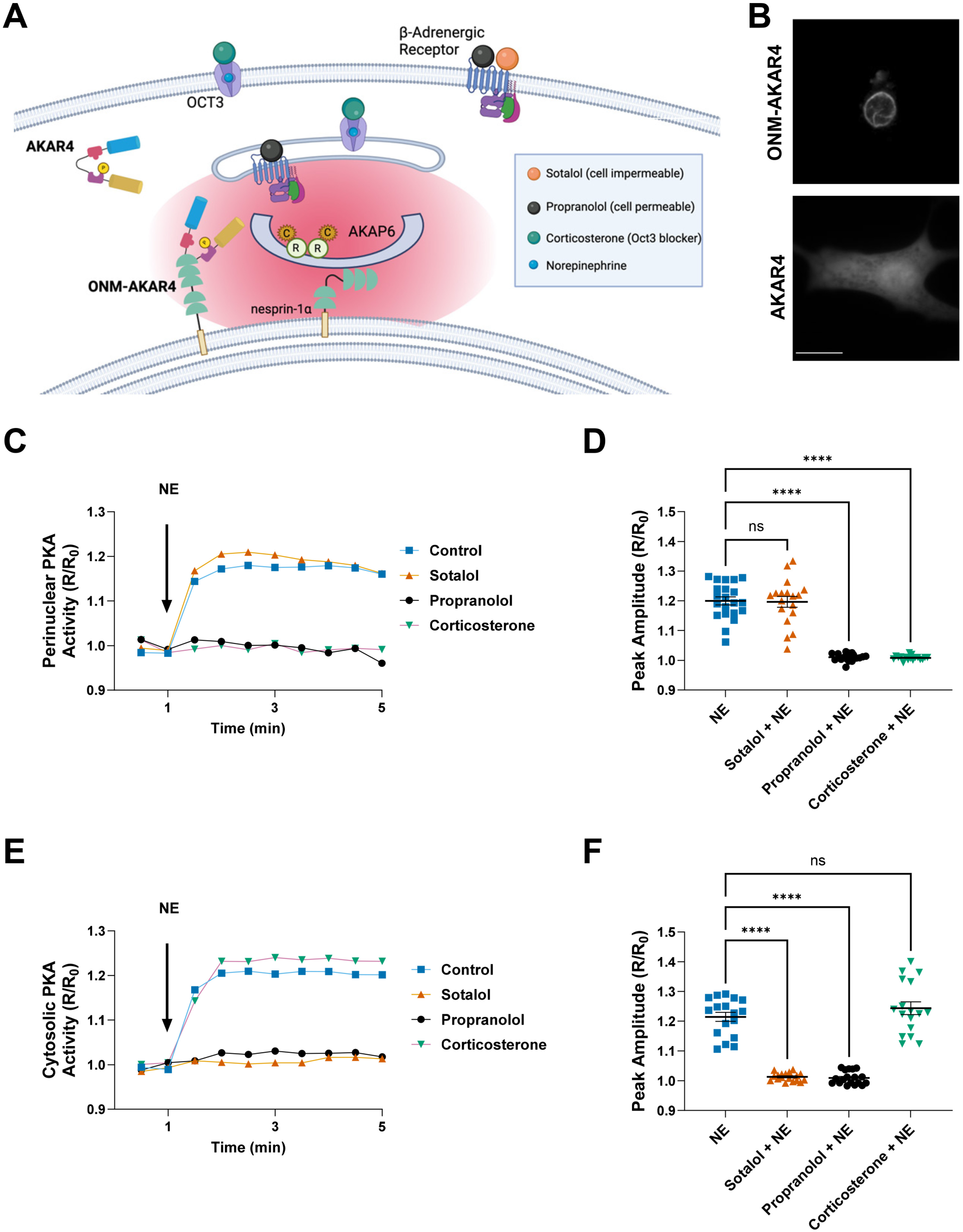
**An intracellular pool of β-adrenergic receptors activates AKAP6β-bound PKA.** (A) Model for experimental design of Figure 1. The AKAR4 FRET biosensor design includes a forkhead-associated domain (FHA1, red) and PKA substrate peptide (LRRATLVD, purple) between Cerulean and cpVenus fluorescent proteins.^67^ AKAR4 phosphorylation increases FRET signal. ONM-AKAR4 is an outer nuclear membrane AKAR4 sensor in which AKAR4 is fused to nesprin-1α, which contains spectrin-like repeat domains (green semicircles) and a C-terminal transmembrane Klarsicht, ANC-1, Syne Homology (KASH) domain (beige).^28^ AKAP6 binds PKA holoenzyme composed of 2 regulatory (R) and 2 catalytic subunits (C). (B) Grayscale images of neonatal rat ventricular myocytes expressing AKAR4 or ONM-AKAR4 (cerulean channel). Bar - 20 µm. (C-F) FRET imaging of norepinephrine (NE)-treated neonatal myocytes expressing AKAR4 (C-D) or ONM-AKAR4 (E-F), following pre-treatment with the indicated inhibitors. Representative tracings and peak amplitudes for FRET ratio (R normalized to baseline R_0_) are shown.

To investigate if AKAP6β-bound PKA is activated by intracellular βARs, we initially used a pharmacologic approach based upon the different membrane permeabilities of βAR antagonists (Fig. 1A). In primary neonatal rat ventricular myocytes, 10 nM norepinephrine (NE) induced a similar increase (∼20%) in biosensor activity whether detected by AKAR4 or ONM-AKAR4 in the cytosol and at the nuclear envelope, respectively (Figure 1C-F). Likewise, NE-stimulated PKA activity was inhibited by the membrane permeable β-blocker propranolol regardless of the location of the FRET biosensor. In contrast, the membrane impermeable β-blocker sotalol inhibited NE-stimulated cytosolic AKAR4, but not perinuclear ONM-AKAR4 PKA signals. These results suggest that in contrast to PKA in the cytosol, the perinuclear AKAP6β compartment is not activated by βARs on the plasma membrane exposed to the outside of the cell.

While βARs on the plasma membrane can be activated by extracellular ligands, local activation of intracellular βARs presumably requires ligand entry into the cell. The endogenous catecholamines norepinephrine and epinephrine can enter cells through the non-selective organic cation transporter-3 (OCT3).^29^ Consistent with the inhibition of cytosolic AKAR4 signals by the membrane impermeant β-blocker sotalol, the OCT3 inhibitor corticosterone^30^ had no effect on NE-induced AKAR4 transients in neonatal myocytes (Figure 1E,F). In contrast, corticosterone completely inhibited NE-induced ONM-AKAR4 signals, which were resistant to sotalol inhibition (Figure 1C,D). Together, these results suggest that PKA confined to the perinuclear AKAP6 signalosome is activated by intracellular βARs binding ligand within the cell, while cytosolic PKA is stimulated by plasma membrane receptors binding ligand outside of the cell. Notably, despite the well-established differences in ultrastructure between cultured primary neonatal and adult rat ventricular myocytes,^31^ similar results were obtained for the β-blockers and corticosterone in experiments employing adult myocytes, with the exception that corticosterone partially inhibited NE-induced cytosolic AKAR4 signals in adult myocytes (Figure S1).

### Perinuclear β-adrenergic receptors are necessary and sufficient for activation of AKAP6β-bound PKA

The camelid single chain nanobody Nb80 binds agonist-occupied β_1_- and β_2_-adrenergic receptors, inhibiting downstream signaling, including cAMP production.^32,33^ To directly target β-adrenergic receptors near AKAP6 signalosomes, we expressed Nb80 in fusion to nesprin-1α and the fluorescent tag mCherry (ONM-Nb80), restricting localization of the nanobody to the ONM (Figure 2A,B). When compared to control mCherry-tagged nesprin-1α (ONM-Control), ONM-Nb80 expression completely inhibited NE-induced PKA perinuclear activity in neonatal myocytes and had a small but significant inhibition (27%) of the subsequent acute activation of cytosolic PKA activity (Figure 2C-F). In adult myocytes, ONM-Nb80 expression had no significant effect on cytosolic PKA, while again completely inhibiting perinuclear PKA activity (Figure S2A-D). These results suggest that active βAR in the vicinity of the ONM is required for activation of AKAP6β-bound PKA.

**Figure 2:**
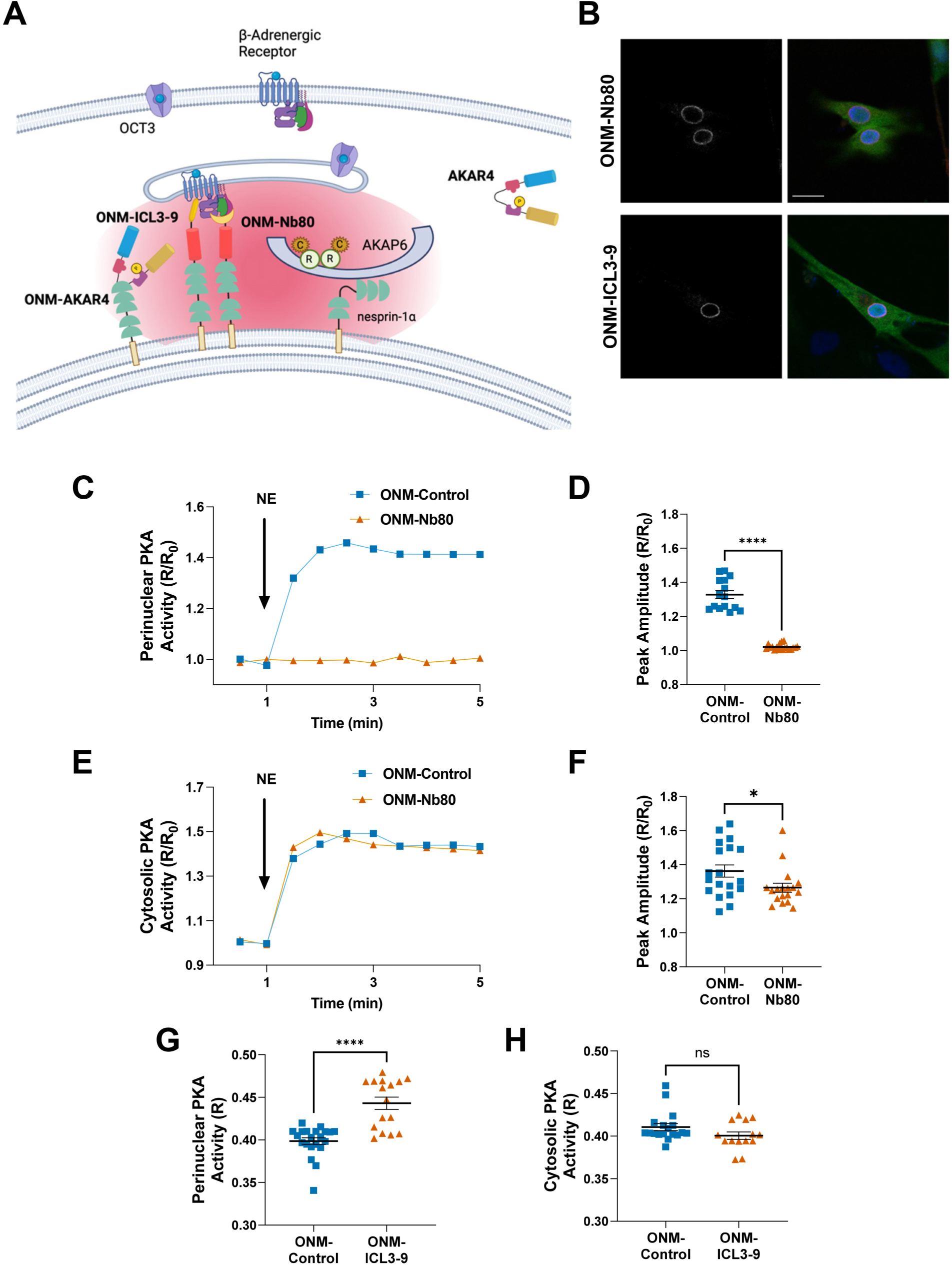
Perinuclear β-adrenergic receptors are necessary and sufficient for activation of AKAP6β-bound PKA. (A) ONM-Nb80 consists of Nb80 nanobody in fusion to nesprin-1α and mCherry, conferring inhibition of βAR downstream signaling. ONM-ICL3-9 contains ICL3-9 pepducin in fusion to nesprin-1α and mCherry, conferring activation of βAR downstream signaling. ONM-control is a nesprin-1α-mCherry fusion protein. (B) Neonatal myocytes expressing AKAR4 (green) and either ONM-Nb80 or ONM-ICL3-9 (red and grayscale images). Bar - 20 µm. (C-F) FRET imaging of NE-treated neonatal myocytes expressing AKAR4 or ONM-AKAR4 and ONM-Control or ONM-Nb80. Representative tracings and peak amplitudes for FRET ratio (R normalized to baseline R_0_) are shown. (G-H) FRET imaging of PKA activity (R) in neonatal myocytes expressing AKAR4 or ONM-AKAR4 and ONM-Control or ONM-ICL3-9.

While Nb80 inhibits agonist-dependent βAR signaling, the pepducin ICL3-9, a peptide based upon the β_2_AR third intracellular loop, selectively activates βAR-dependent G_αs_ signaling in the absence of ligand binding and β-arrestin recruitment.^34^ In order to test whether βAR activation at or near the ONM is sufficient for activation of AKAP6β signalosomes, ICL3-9 was similarly expressed in fusion to nesprin-1α and mCherry (ONM-ICL3-9, Figure 2B,G,H). As expected, perinuclearly restricted expression of the pepducin had no effect on steady-state PKA activity in unstimulated neonatal myocytes detected by the cytosolic AKAR4 parent sensor. Remarkably, expression of ONM-ICL3-9 increased ONM-AKAR4 FRET ratio in the absence of any ligand stimulus when compared with ONM-control, implying direct activation of βARs near AKAP6β signalosomes on the ONM. Similar results for ONM-Nb80 and ONM-ICL3-9 were obtained using adult myocytes (Figure S2). Thus, consistent with the results obtained using βAR and OCT3 inhibitors, expression of the Nb80 and ICL3-9 mcherry-nesprin-1α fusion proteins demonstrated that activation of βAR at or near the ONM is necessary and sufficient for activation of PKA at AKAP6β signalosomes, thereby showing the functional relevance of intracellular βARs to compartment-specific signal transduction.

### Golgi-localized βARs activate PKA in AKAP6β signalosomes

Besides the canonical plasma membrane location of GPCRs, βARs have been detected on multiple internal membranes including endosomes, Golgi apparatus, and the nuclear envelope, both as part of receptor recycling and down-regulation, as well as mediating downstream signaling within the cell.^5–8^ It is, therefore, not obvious on which membranes are present the βARs activating AKAP6β signalosomes. For example, in cardiac myocytes the Golgi is in close proximity to the ONM, such that the Golgi and ONM are bridged by nesprin-1α – AKAP6β - AKAP9 – GM130 protein complexes (Figure 3A).^35^ Deep segments of the transverse tubule system, which is an invagination of the plasma membrane, have also been found near the nuclear envelope,^36^ albeit the aforementioned pharmacological data (Figure 1) do not support a role for βARs exposed to the outside of the myocyte in the activation of AKAP6β signalosomes. To determine which organelles might house βARs responsible for activation of PKA at AKAP6β and nesprin-1α, in addition to nesprin-1α fusion targeting to the ONM, Nb80 was targeted to the Golgi by fusion to a beta-1,4-galactosyltransferase 1 targeting sequence (GalT aa 2-79, “Golgi-Nb80”),^7^ to endosomes by fusion to tandem FYVE domains from hepatocyte growth factor 1 - regulated tyrosine kinase substrate (Hrs aa 147-223, “Endo-Nb80”),^37^ and to the plasma membrane by fusion to the N-terminal domain of AKAP7α (aa 1-25, “PM-Nb80”, Figure 3B).^38^ In each case Nb80 was directed to the cytosolic face of these organelles. βARs have also been detected on the sarcoplasmic reticulum, which contains sections that are in close proximity to the ONM.^6,36^ We were unable to direct Nb80 solely to the sarcoplasmic reticulum, as the ONM contains similar proteins as the sarcoplasmic reticulum, with which it is contiguous.^39^

**Figure 3.**
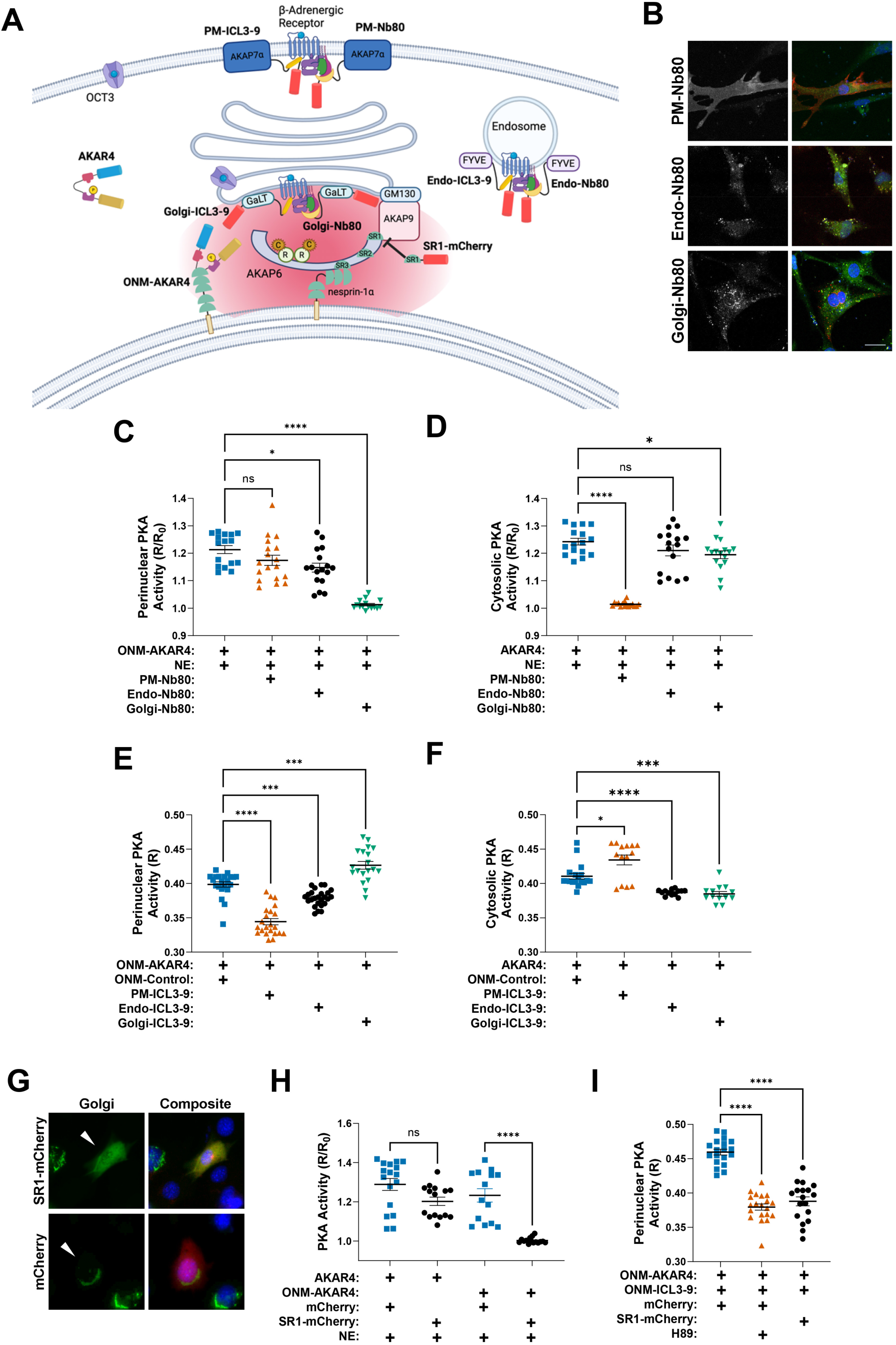
Golgi-localized βARs activate PKA in AKAP6β signalosomes. (A) ICL3-9 and Nb80 fusion peptides containing a mCherry tag were targeted to the plasma membrane (PM), endosome (Endo), or Golgi apparatus (Golgi). SR-1-mCherry contains the first spectrin repeat of AKAP6β that binds AKAP9 and competes their binding. (B) Neonatal myocytes stained with Dapi (blue) and expressing AKAR4 (green) and either PM-Nb80, Endo-Nb80, or Golgi-Nb80 (red and grayscale). Bar - 20 µm. (C-D) FRET imaging of neonatal myocytes expressing AKAR4 or ONM-AKAR4 and either PM-Nb80, Endo-Nb80 or Golgi-Nb80. Fold response to NE is indicated (R normalized to baseline R_0_). See also Figure S3 A,B. (E-F) FRET ratios (R) for neonatal myocytes expressing AKAR4 or ONM-AKAR4 and either OMN-Control, PM-ICL3-9, Endo-ICL3-9, or Golgi-ICL3-9. Note that the data for OMN-Control are duplicated from Figure 2G-H, as panels E and F were performed as an extension of the same study. (G) Neonatal myocytes expressing SR-1-mCherry or mCherry (red, cells with arrowheads), GalT-GFP (Golgi, green) and stained with Dapi (blue). (H) FRET imaging of myocytes expressing AKAR4 or ONM-AKAR4 and either mCherry or SR1-mCherry. and treated with NE as indicated. Fold response to NE is indicated (R normalized to baseline R_0_). See also Figure S3H. (I) FRET ratios (R) for neonatal myocytes expressing ONM-AKAR4 and either mCherry or SR1-mCherry and/or treated with the PKA inhibitor H89.

Consistent with the effects of sotalol and corticosterone suggesting that the cytosolic PKA compartment is strongly regulated by extracellular agonist stimulation of βARs, expression of Nb80 on the plasma membrane of neonatal myocytes completely inhibited the NE-induction of PKA activity in the cytosol, while having no significant effect on NE-stimulated perinuclear PKA activity detected with ONM-AKAR4 (Figure 3C&D and Figure S3A). Expression of Nb80 on endosomes did not significantly affect NE-stimulated cytosolic PKA activity, while partially (31%) inhibiting NE-stimulated perinuclear PKA activity. In contrast, localization of Nb80 to Golgi completely blocked the NE activation of perinuclear PKA as detected by ONM-AKAR4, while conferring a small (19%) but significant inhibition of NE-stimulated cytosolic PKA activity. Taken together with results for ONM-Nb80, these findings suggest that βARs within a compartment in the vicinity of both the Golgi and ONM are primarily responsible for activation of PKA at AKAP6β signalosomes, while βARs on the plasma membrane are primarily responsible for activation of cytosolic PKA.

To corroborate these results, the βAR-activating pepducin ICL3-9 was similarly targeted to the plasma membrane, endosomes, and Golgi. Steady-state cytosolic PKA activity detected with the parent AKAR4 sensor was increased 6% in unstimulated myocytes constitutively expressing PM-ICL3-9, while, resulting in a 14% decrease in ONM-AKAR4 FRET ratio (Figure 3E&F). Targeting of the pepducin to endosomes inhibited cytosolic and perinuclear PKA activity 5% and 6%, respectively. Notably, Golgi-ICL3-9 increased steady-state perinuclear PKA activity in myocytes detected with ONM-AKAR4 7%, while like Endo-ICL3-9 inhibiting signals detected with the cytosolic parent sensor 6%. Taken together with the aforementioned results, these findings suggest that βAR within the vicinity of the Golgi and ONM are necessary and sufficient for activation of PKA in the AKAP6β compartment.

The above results place the βAR activating AKAP6β-bound PKA in a perinuclear compartment. To distinguish between βARs that are on the ONM and the Golgi, we took advantage of the prior observation that the Golgi can be separated from the nuclear envelope by disrupting the nesprin-1α – AKAP6 - AKAP9 – GM130 protein bridge between the two organelles.^35^ While the third spectrin repeat domain (SR3) of AKAP6 binds nesprin-1α,^13^ the first AKAP6 spectrin repeat domain (SR1) binds AKAP9, such that expression of an SR1-mCherry fusion protein will compete AKAP6-AKAP9 binding releasing the Golgi from its perinuclear location.^35^ As shown in Figure 3G, Golgi, distinctly labelled by expression of a GalT-GFP fusion protein in control mCherry expressing cells, was dispersed in myocytes expressing SR1-mCherry. As expected, SR1-mCherry expression had no significant effect on AKAR4 signals induced by NE (Figure 3H and Figure S3C). Notably, SR1-mCherry expression completely suppressed NE-stimulated perinuclear PKA activity detected with ONM-AKAR4. In addition, SR1-mCherry-mediated Golgi dispersion completely inhibited perinuclear PKA elicited by ONM-ICL3-9 just like pharmacological PKA inhibition with H89 (Figure 3I, compare to Figure 3E). These data suggest that βAR on the Golgi close enough to the ONM to be regulated by ICL3-9 and Nb80 localized to either membrane is responsible for activating cAMP signaling at AKAP6β signalosomes.

### Defining the dimensions of the AKAP6β cAMP compartment

That single polypeptide βAR activators localized to the ONM or Golgi were able to activate PKA activity detected with ONM-AKAR4 without increasing cytosolic AKAR4 signals suggested that AKAP6β signalosomes are present within a discrete, independent cAMP signaling compartment. To estimate the size of the AKAP6β cAMP compartment, we expressed in neonatal myocytes nesprin-1α fusion sensors containing the ratiometric cAMP FRET sensor Epac2-camps (Figure 4A,B),^40^ one version with (ONM-50-Epac2-camps) and one without (ONM-Epac2-camps) an intervening rigid spacer comprising 50 copies of the pentapeptide EAAAK, which would further distance the sensor from the ONM by ∼10-26 nm.^41^ Just as NE could activate PKA detected with both perinuclear and cytosolic AKAR4 (Figure 1), NE similarly activated ONM-Epac2-camps, ONM-Epac2-50-camps and parental Epac2-camps sensors (Figure 4C). In addition, cAMP levels at AKAP6β signalosomes detected with ONM-Epac2-camps were similarly elevated whether in response to NE or expression of the βAR-activating ONM-ICL3-9 fusion protein (Figure 4D). Remarkably, insertion of the (EAAAK)_50_ spacer completely prevented the detection by the FRET biosensor of perinuclear cAMP induced by the pepducin fusion protein, while, as a control, the longer cAMP sensor was still equally responsive to NE stimulation (Figure 4E). These results imply that cAMP levels at AKAP6β signalosomes drop precipitously from their local source of origin, within a distance at least 2 orders of magnitude smaller than the size of the overall nucleus (∼10 µm in diameter).^13^

**Figure 4.**
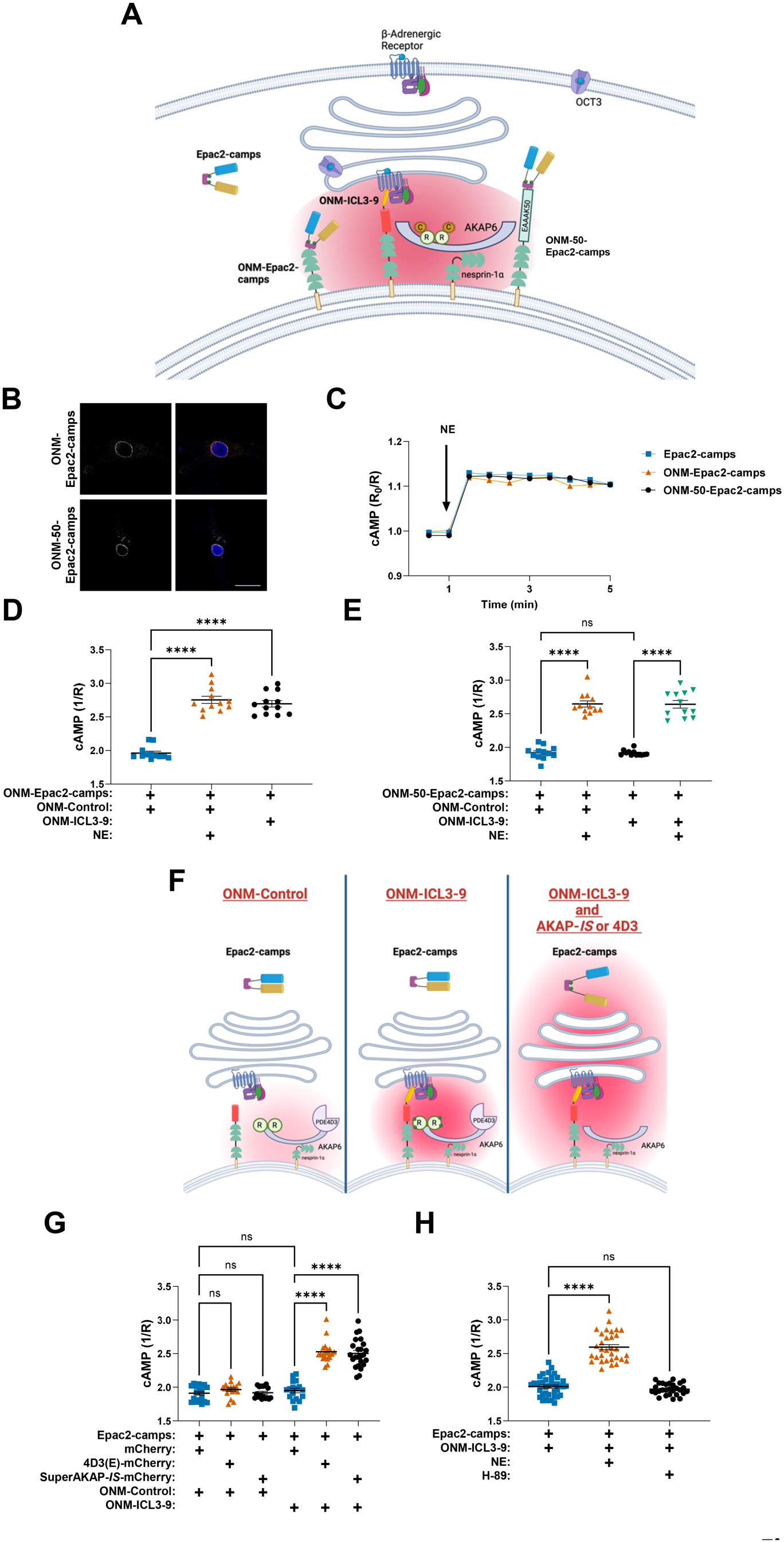
Defining the dimensions of the AKAP6β cAMP compartment. (A) ONM-targeted Epac2-camps cAMP FRET sensor was expressed with (ONM-50-Epac2-camps) and without (ONM-Epac2-camps) an intervening 50 copies of the pentapeptide EAAAK, which increases the distance of sensor from the ONM by ∼10-26 nm. (B) ONM-Epac2-camps and ONM-50-Epac2-camps expression (red and grayscale) in neonatal myocytes stained with Dapi and expressing also ONM-ICL3-9 (green). Bar - 20 µm. _(C)_ Representative tracings of myocytes expressing Epac2-camps, ONM-Epac2-camps or ONM-50-Epac2-camps showing similar responses to NE. The reciprocal of normalized FRET ratios for this biosensor (R_0_/R) are shown as Epac2-camps exhibits decreased FRET following cAMP binding.^40^ (D-E) FRET imaging of neonatal myocytes expressing ONM-Epac2-camps or ONM-50-Epac2-camps and ONM-mCherry or ONM-ICL3-9. Cells were stimulated for 1 day with NE, and cAMP levels are reported as the reciprocal of steady state FRET ratio (1/R). (F) Model for regulation of AKAP6 cAMP compartment. Super-AKAP-IS and 4D3(E)-mCherry compete PKA and PDE4D3 binding to AKAP6, respectively. (G,H) Neonatal myocytes expressing Epac2-cAMPs cytosolic sensor and the indicated recombinant proteins were cultured in the absence or presence of NE and H-89 PKA inhibitor. cAMP levels are reported as the reciprocal of steady state FRET ratio (1/R).

Mechanisms limiting the size of cAMP compartments include physical membrane barriers, the activity of localized phosphodiesterases, and buffering by cAMP-binding proteins.^2^ The results above suggest that AKAP6β signalosomes are in a cleft between the Golgi and ONM, but the proximity of these membranes does not entirely explain the results with Epac-camps fusion sensors. AKAP6β binds the cAMP-specific, PKA-activated phosphodiesterase PDE4D3, that in the absence of local stimulation can minimize cAMP levels and that in the presence of compartment activation can limit cAMP signaling via negative feedback regulation.^19,24^ AKAP6β, as well as potentially AKAP9 in the compartment, also binds PKA holoenzyme containing two RII-subunits that bind cAMP.^25^ If PDE4D3 local cAMP hydrolysis or PKA RII buffering confers cAMP compartmentation, loss of PDE4D3 or PKA association with the signalosome should increase the distance at which cAMP can accumulate to levels capable of mediating signal transduction (Figure 4F). To test this hypothesis, cytosolic cAMP was assayed in myocytes expressing the parental, cytosolic Epac2-camps sensor in the absence or presence of recombinant polypeptides that compete PDE4D3 or PKA binding to AKAP6β. 4D3(E)-mCherry contains a PDE4D3-derived peptide based upon the N-terminal domain of PDE4D3 that binds AKAP6,^28^ while SuperAKAP-*IS*-mCherry contains a synthetic peptide that will compete PKA RII subunit binding to AKAPs.^42^ Consistent with results obtained for AKAR4, expression of perinuclearly localized pepducin (ONM-ICL3-9) had no effect on cAMP levels detected by the cytosolic Epac2-camps parent sensor (Figure 4G). In addition, both PDE4D3 and PKA displacement from AKAP6β by expression of 4D3(E)-mCherry and SuperAKAP-*IS*-mCherry, respectively, had no effect on cytosolic cAMP levels in unstimulated cells expressing ONM-control. Remarkably, expression of either the PKA or PDE4D3 anchoring disruptor peptide in conjunction with activation of perinuclear βARs by ONM-ICL3-9 resulted in cytosolic accumulation of cAMP sufficient to activate Epac2-camps. To distinguish PKA effects on cAMP buffering from PDE4D3 feedback activation, ONM-ICL3-9 was also expressed in the presence of the PKA inhibitor H89, but in the absence of any anchoring disruptor peptide, resulting in no activation of cytosolic Epac2-cAMP (Figure 4H). As inhibition of PKA anchoring by SuperAKAP-*IS*-mCherry, but not inhibition of PKA activity by H89 resulted in spread of cAMP into the cytosolic compartment, we suggest that cAMP buffering by PKA RII-subunits, in conjunction with PDE4D3 local cAMP hydrolysis, restricts locally generated cAMP to the AKAP6β compartment, thereby conferring independence of this small compartment from cAMP signaling elsewhere in the cell.

### Regulation of Ca^2+^-dependent calcineurin signaling by AKAP6β-associated βARs

The activation of cAMP signaling at AKAP6β signalosomes by local βARs would presumably entail the activation by downstream signaling known to be dependent upon perinuclear cAMP. Calcineurin is a Ca^2+^/calmodulin-dependent phosphatase that activates NFAT and is required for pathological cardiac hypertrophy.^11^ We recently showed that in addition to orchestrating perinuclear cAMP signaling, AKAP6β organizes a perinuclear Ca^2+^ signaling compartment that is independent of Ca^2+^ cycling regulating excitation-contraction coupling, but is required for local activation of the calcineurin-NFAT pathway and induction of myocyte hypertrophy.^27^ In particular, perinuclear Ca^2+^ - calcineurin signaling was dependent upon AKAP6β expression and elevated local cAMP levels. The effects of local cAMP were apparently through PKA-catalyzed phosphorylation of associated ryanodine receptors, which could release Ca^2+^ from intracellular stores into the perinuclear compartment. Thus, in concert with the above results, we hypothesize that Golgi βARs activating AKAP6β signalosomes serve to regulate local pro-hypertrophic Ca^2+^ - calcineurin signaling.

To test whether perinuclear βARs regulate the perinuclear Ca^2+^ signaling compartment, we first expressed in neonatal myocytes a nesprin-1α fusion protein containing the intensiometric Ca^2+^ sensor GCaMP6s (ONM-GCaMP6s, Figure 5A,B).^43^ Consistent with the inhibition of perinuclear PKA activity (Figure 2D), inhibition of perinuclear βARs with ONM-Nb80 prevented the observed persistent elevation in perinuclear [Ca^2+^] in response to chronic βAR stimulation, while having no effect on cytosolic [Ca^2+^] assayed using the diffusely localized, parent GCaMP6s sensor (Figure 5C,D). In addition, consistent with the activation of perinuclear βARs and PKA (Figure 2G), ONM-ICL3-9 expression increased steady-state perinuclear [Ca^2+^] in unstimulated cells, while not affecting cytosolic [Ca^2+^] measured with the parent GCaMP6s sensor (Figure 5E,F). Further, consistent with our previously published findings that Ca^2+^ influx was dependent upon PKA activity,^27^ ONM-ICL3-9 activation of ONM-GCaMP6s was inhibited by H89. These results show that βARs associated with AKAP6β signalosomes not only regulate local PKA activity, but also Ca^2+^ fluxes within that compartment.

**Figure 5.**
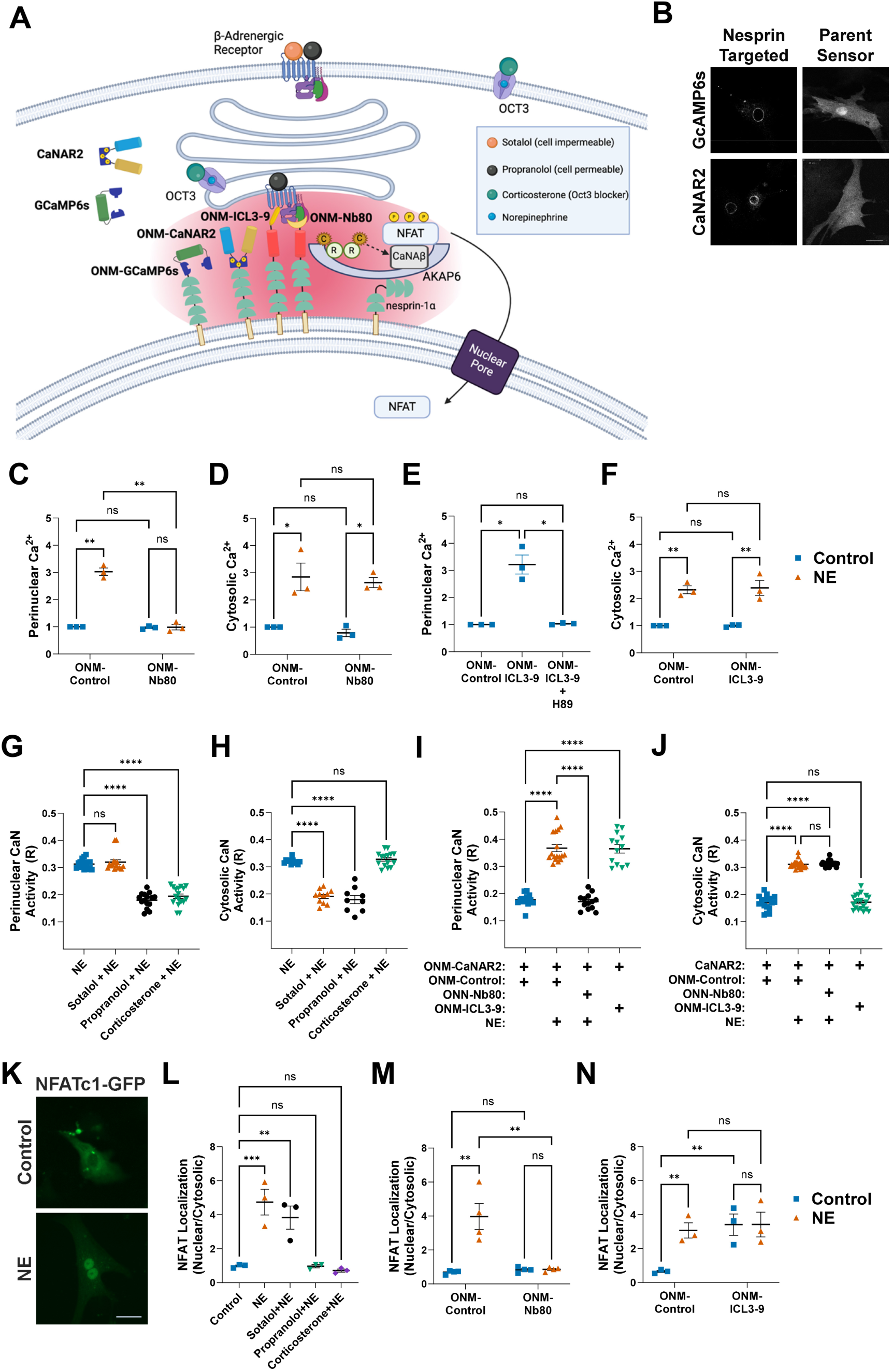
Regulation of Ca^2+^-dependent calcineurin signaling by AKAP6β-associated βARs. (A) CaNAR2 is a Cerulean 3 - YPet FRET biosensor for CaN activity based upon the dephosphorylation of a NFATc2 fragment.^45^ GCaMP6s is a circularly permuted green fluorescent protein (cpGFP) intensiometric Ca^2+^ sensor based upon binding of the M13 peptide to a variant calmodulin.^43^ (B) Images are shown in grayscale of neonatal myocytes expressing cytosolic and nesprin-targeted (ONM) GCaMP6s and CaNAR2. Bar – 20 µm. (C-F) Ca^2+^ levels were measured in neonatal myocytes expressing ONM-Control, ONM-Nb80 or ONM-ICL3-9 and stimulated for 1 day with NE. Ca^2+^ levels are expressed as average fold sensor fluorescence for cells acquired for each of 3 independent preparations. (G-J) CaN activity was measured in neonatal myocytes expressing ONM-Control, ONM-Nb80 or ONM-ICL3-9 and treated for 1 day with NE and the indicated inhibitors. CaN activity is reported as FRET ratios (R). (K) Neonatal myocytes expressing NFATc1-GFP and stained with Flag tag antibody. Examples are shown of cells with mainly cytosolic or nuclear NFATc1 localization. Bar – 20 µm. (L-M) NFATc1-GFP localization in myocytes expressing ONM-localized fusion proteins and treated for 1 day with NE and the indicated inhibitors. Average relative localization is reported for cells acquired for each of 3-4 independent preparations.

Calcineurin is recruited during prolonged βAR stimulation to AKAP6β signalosomes, where elevated local [Ca^2+^] promotes calcineurin-NFAT signaling driving myocyte hypertrophy.^27,44^ To test whether elevated perinuclear PKA activity and [Ca^2+^] in response to chronic stimulation of perinuclear βARs was associated with activation of the calcineurin-NFAT pathway, we imaged myocytes expressing a nesprin-1α-localized version of the calcineurin FRET biosensor CaNAR2 (ONM-CaNAR2, Figure 5A,B).^27,45^ Myocytes were studied as above using similar pharmacologic and molecular approaches. Like ONM-AKAR4, propranolol, but not sotalol inhibited NE-induced perinuclear calcineurin activity, while, like AKAR4, both β-blockers inhibited NE-induced cytosolic calcineurin activity detected with the parental CaNAR2 sensor (Figure 5G,H). Notably, the OCT3 inhibitor corticosterone only inhibited βAR-induced calcineurin activity detected with the perinuclear ONM-CaNAR2 sensor, confirming a role for intracellular βARs in the regulation of perinuclear calcineurin signaling. Expression of ONM-Nb80, which inhibits perinuclear βARs, selectively inhibited NE-stimulated perinuclear calcineurin activity (Figure 5I,J). Conversely, expression of ONM-ICL3-9, which activates perinuclear βARs, induced perinuclear calcineurin activity, but not calcineurin activity detected with the parent CaNAR2 sensor. Similar results were obtained for CaN signaling in adult myocytes (Figure S4).

When dephosphorylated by calcineurin phosphatase, NFAT transcription factor translocates into the myocyte nucleus promoting hypertrophic gene expression.^11^ As expected, the nuclear accumulation of GFP-tagged NFATc1 in neonatal myocytes chronically treated with NE was inhibited by the cell permeable β-blocker propranolol (Figure 5K-L). NE-induced GFP-NFATc1 nuclear translocation was not inhibited by membrane impermeant sotalol, but the OCT3 inhibitor corticosterone completely blocked nuclear translocation. Consistent with the regulation of perinuclear calcineurin, ONM-Nb80 inhibited NE-induced GFP-NFATc1 nuclear translocation, while ONM-ICL3-9 induced GFP-NFATc1 nuclear translocation in the absence of agonist (Figure 5M,N). Taken together, these results suggest that perinuclear βARs, which regulate cAMP-PKA signaling in the AKAP6β compartment, similarly regulate perinuclear [Ca^2+^] and the activity of the calcineurin – NFAT pathway.

### Regulation of myocyte hypertrophy by perinuclear β-adrenergic receptors

AKAP6β signalosomes, including AKAP6β-bound PKA and calcineurin, have been shown to be required for myocyte hypertrophy (non-mitotic cell growth).^21,44^ The regulation of AKAP6β signalosomes by perinuclear βARs implies a unique role for these compartment-specific GPCRs in the regulation of myocyte hypertrophy. Culture of primary adult myocytes in the presence of NE results in an increase in length and width of the relatively columnar-shaped cells.^27^ To test directly whether perinuclear βARs regulate myocyte hypertrophy, adult myocytes chronically stimulated with NE for 2 days were treated as above to alter perinuclear βAR activity. Treatment with propranolol and corticosterone, but not sotalol, prevented the NE-induced myocyte growth in length and width (Figure 6A-F, Figure S5A). In addition, expression of ONM-Nb80 inhibited NE-induced myocyte hypertrophy, while expression of ONM-ICL3-9 induced myocyte growth in length and width independently of GCPR stimulation. These results imply that activation of perinuclear βARs is necessary and sufficient for the induction of adult myocyte hypertrophy by AKAP6β signalosomes.

**Figure 6:**
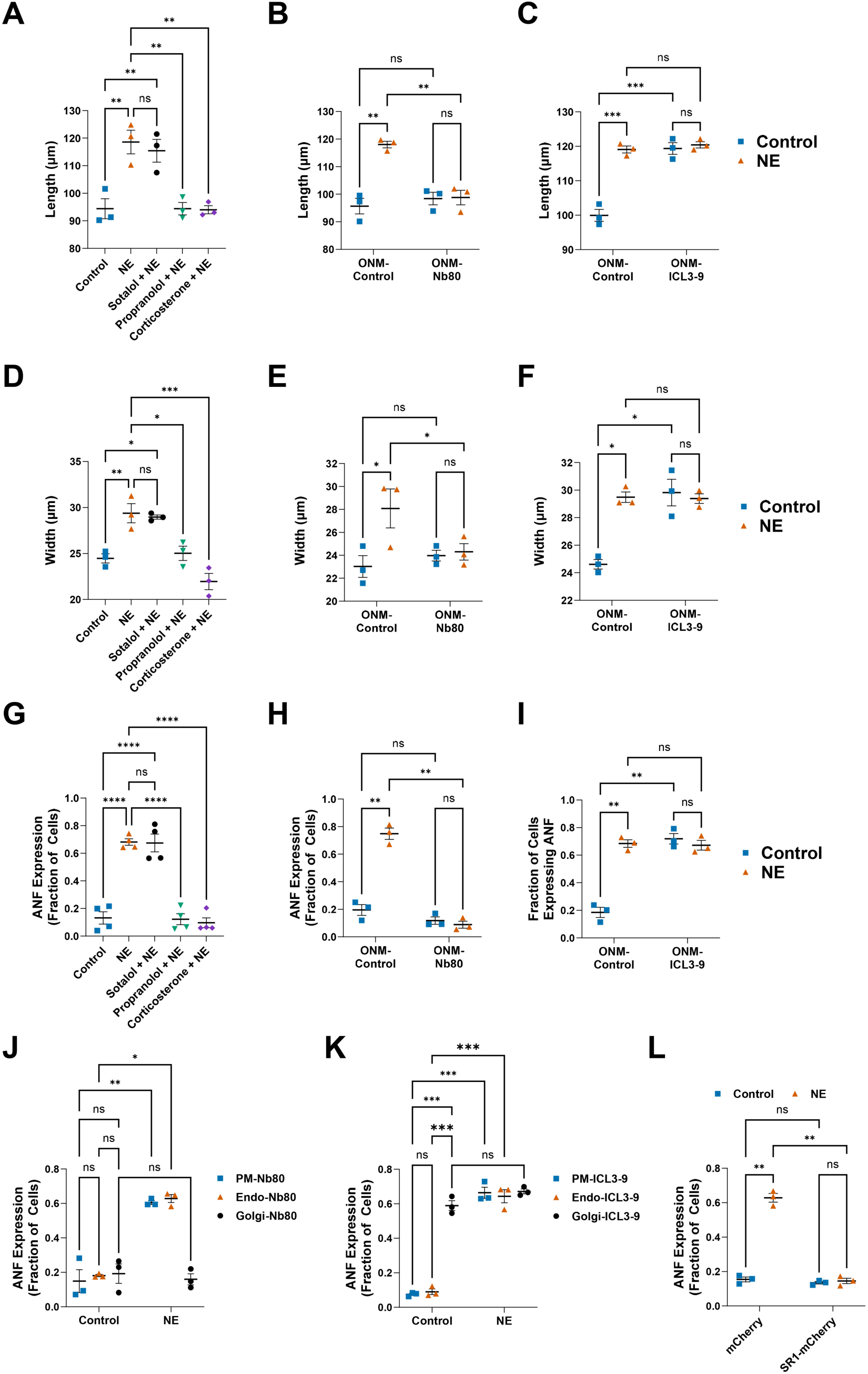
Regulation of myocyte hypertrophy by perinuclear β-adrenergic receptors. (A-F) Adult rat ventricular myocytes RNV were treated for 2 days with either no drug (Control), NE or the indicators inhibitors and/or infected with adenovirus to express ONM-Control, ONM-Nb80, or ONM-ICL3-9 before morphometric analysis for myocyte length and width. Datapoints represent the mean values of >50 cells measured for each of 3 biological replicates. (G-L) Neonatal rat ventricular myocytes expressing ONM-Control, ONM-Nb80, ONM-ICL3-9, PM-Nb80, Endo-Nb80, Golgi-Nb80, PM-ICL3-9, Endo-ICL3-9, Golgi-ICL3-9, SR1-mCherry, or mCherry control were treated for 2 days with either no drug (Control), NE or the indicators inhibitors before staining for ANF. Datapoints represent the fraction of cells with prominent ANF staining for >50 cells measured for each of 3 biological replicates. See also Figure S5.

In contrast to other AKAP scaffolds, AKAP6β expression is not cardioprotective, while required for pathological cardiac remodeling in response to diverse pathophysiological insults.^22,23^ Symmetric growth in length and width of adult myocytes may represent physiological or pathological hypertrophy. However, elevated atrial natriuretic factor (ANF) expression is a marker for pathological myocyte hypertrophy readily detected in neonatal myocytes stimulated with GPCR agonists.^46^ Consistent with the above results, NE-induced ANF expression in neonatal myocytes was inhibited by propranolol and corticosterone, but not sotalol (Figure 6G). In addition, NE-induced ANF expression was suppressed by ONM-Nb80, that directly inhibits perinuclear βAR, while ANF expression was induced independently of NE stimulation by ONM-ICL3-9, that directly activates perinuclear βAR (Figure 6H,I). Importantly, this perinuclear pool of βAR is both necessary and sufficient to drive pathological gene transcription, as localization of Nb80 to the Golgi, but not the plasma membrane or endosomes, inhibited NE-induced ANF expression. Conversely, ICL3-9 pepducin localization to the Golgi, but not the plasma membrane or endosomes, resulted in agonist-independent ANF expression (Figure 6J,K). Finally, expression of SR1-mCherry, that disrupts AKAP6-AKAP9 binding, resulting in Golgi dispersion, also inhibited NE-induced ANF expression (Figure 6L). Taken together, these results suggest that in contrast to plasma membrane βAR stimulation activating PKA elsewhere in the myocyte, Golgi-localized βARs regulate perinuclear AKAP6β signalosomes responsible for the coordinated activation of cAMP-and Ca^2+^-dependent signaling pathways inducing pathological myocyte hypertrophy.

### Perinuclear β-adrenergic receptors regulate pathological cardiac remodeling in vivo

The regulation of pathological myocyte hypertrophy by perinuclear βARs *in vitro* suggested that this compartmentalized GPCR pool would be similarly critical for the response of the heart in disease. To demonstrate that GPCRs in a privileged intracellular compartment is required for the determination of organ phenotype *in vivo*, we expressed ONM-Nb80 and ONM-ICL3-9 in mice using adeno-associated virus (AAV) gene therapy vectors. AAV serotype 9 vectors were generated that express under the control of the cardiac myocyte-specific cardiac troponin T (cTnT) promoter the ONM-Nb80 and ONM-ICL3-9 constructs, in this case without the mCherry tag due to the limited length (∼4.8 kb) of AAV genomes (Figure 7A).^47^ When administered intravenously to adult mice, these vectors expressed ONM-Nb80 and ONM-control fusion proteins at lower levels in the heart than endogenous nesprin-1α (Figure S6). ONM-ICL3-9 was expressed at even lower levels and was not consistently detected by western blot, but by qRT-PCR (data not shown).

**Figure 7.**
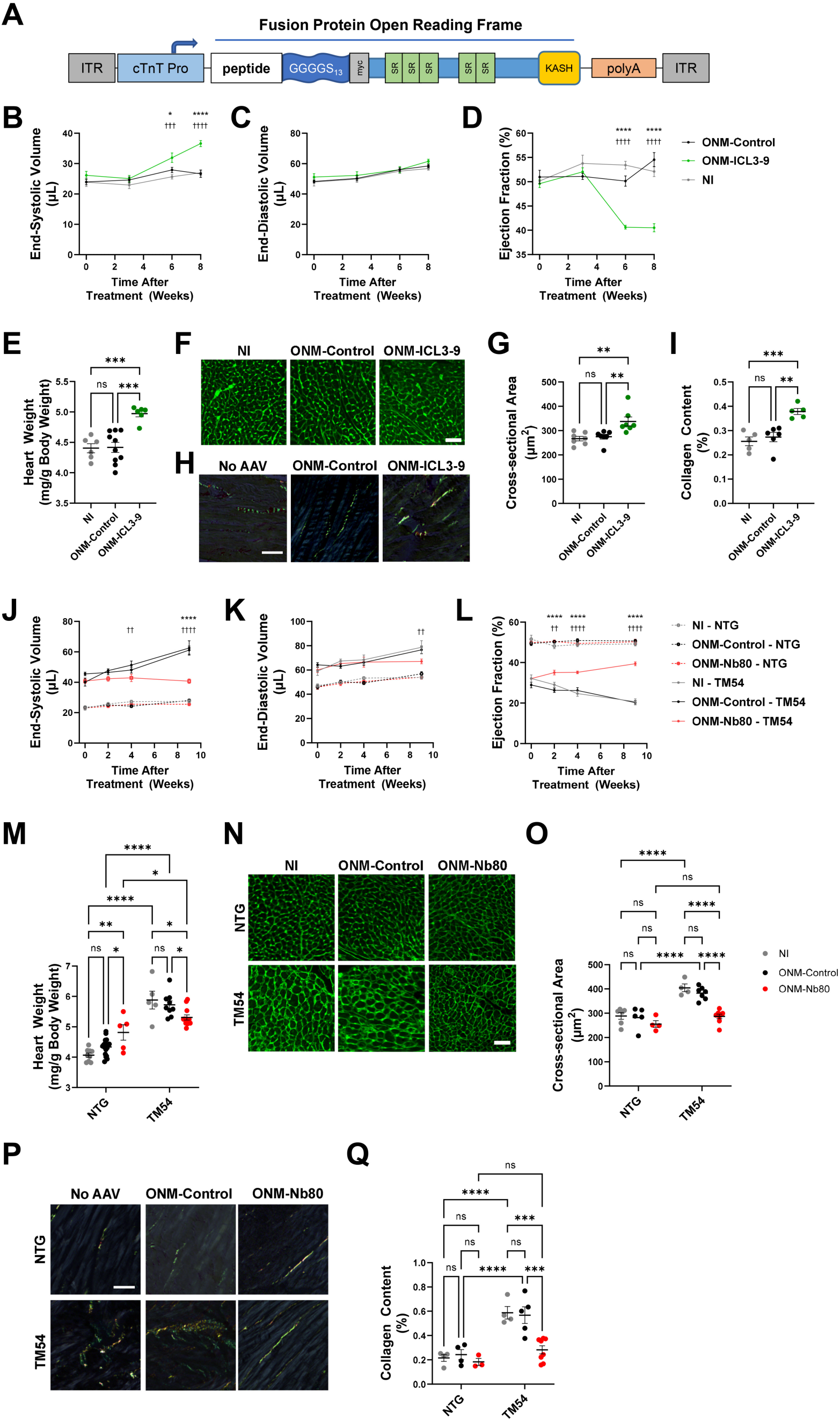
Perinuclear β-adrenergic receptors regulate pathological cardiac remodeling in vivo. (A) Cardiomyocyte-selective AAV9 vectors containing a cTnT promoter were used to express versions of ONM-Control, ONM-ICL3-9, and ONM-Nb80 lacking the mCherry tag. (B-D) C57BL6/NJ mice treated at 6-8 weeks of age with 5×10^11^ vg i.p AAV9 or no virus (NI) were followed by serial “4D” echocardiography for 8 weeks. *n* = 6-9. * vs. ONM-Control, ^†^ vs. NI controls. See Videos S1-S3. (E) Indexed heart weight determined gravimetrically at 8 week endpoint. (F,G) Myocyte cross-sectional area at endpoint determined by wheat-germ agglutinin staining and fluorescent microscopy. (H,I) Interstitial myocardial fibrosis at endpoint determined by Picrosirius Red staining and polarized light microscopy. (J-L) FVB/N TM54 and non-transgenic (NTG) littermate mice were mice treated with AAV as above and followed by serial “4D” echocardiography for 8 weeks. *n* = 8-15. * vs. ONM-Control, ^†^ vs. NI controls. All corresponding TM54 and NTG cohorts were significantly different at all time points (*p* < 0.01). See Videos S4-S9. (M) Indexed heart weight determined gravimetrically at 10 week endpoint. (N,O) Myocyte cross-sectional area at endpoint determined by wheat-germ agglutinin staining and fluorescent microscopy. (P,Q) Interstitial myocardial fibrosis at endpoint determined by Picrosirius Red staining and polarized light microscopy.

Expression of ONM-ICL3-9 in wildtype adult C57BL/6NJ mice resulted in the development of prominent systolic cardiac dysfunction within 2 months of AAV administration, as detected by “4D” echocardiography (Figure 7B-D and Video S1-3). Left ventricular (LV) end-systolic volume was increased, while LV ejection fraction was decreased significantly by 6 weeks after AAV injection in comparison to ONM-nesprin control or non-treated naive mice, which had preserved cardiac function. Although the hearts were not significantly dilated in diastole following 2 months of ONM-ICL3-9 expression, by gravimetric analysis the hearts exhibited significant ventricular hypertrophy (13% increased indexed heart weight, Figure 7E). Likewise, cardiac myocyte cross-section area measured in wheat-germ agglutinin-stained LV tissue sections was increased in mice expressing ONM-ICL3-9, when compared to the two control cohorts (Figure 7F,G). Notably, indicative of pathological remodeling, an increase in interstitial myocardial fibrosis was detected following ONM-ICL3-9 expression (Figure 7H,I). Taken together, these results demonstrated that, like *in vitro,* activation of perinuclear βAR is sufficient to induce cardiomyopathy *in vivo*.

To test whether perinuclear βARs are required for pathological cardiac remodeling, we conducted a treatment study in a mouse model of human Familial Dilated Cardiomyopathy. The “TM54” FVB/N mouse contains a transgene that expresses under the control of the cardiac myocyte-specific α-myosin heavy chain promoter a mutant cDNA for the sarcomeric protein α-tropomyosin.^48^ The α-tropomyosin E54K mutation causes decreased myofilament Ca^2+^ sensitivity and tension development, which manifests as prominent cardiac systolic dysfunction and early onset cardiomyopathy.^48^ 6-8-week-old TM54 mutant mice with obvious cardiac dysfunction and non-transgenic (NTG) littermate mice with normal function were randomized for treatment with AAV to express ONM-Nb80 or ONM-control or for no treatment at all. During the 9-weeks following treatment, ONM-control and non-injected AAV-naive TM54 mice exhibited a progressive decline in cardiac systolic function, as evident by a significant decline in LV ejection fraction (11.5% and 8.7% decrease in LV ejection fraction, respectively, for non-injected and ONM-control cohorts, respectively, *p* < 0.0001 vs. initial time point, Figure 7L). Remarkably, TM54 mice treated with ONM-Nb80 exhibited an improvement over the 10-week study of 7.3% in ejection fraction (*p* < 0.0001 vs. initial time point), resulting in a 19% improvement in ejection fraction at endpoint in comparison to the control cohorts (*p* < 0.0001). The improvement in function was also evident by the significantly less LV end-systolic volume for OMN-Nb80-treated mice than control mice (33% and 35% lower than ONM-Control and non-injected mice, respectively, *p* < 0.0001, Figure 7J). Notably, neither ONM-Nb80 nor ONM-Control had any significant effect on the cardiac function of wildtype littermates. The improvement in TM54 cardiac function conferred by ONM-Nb80 treatment was accompanied by decreased cardiac hypertrophy (Figure 7M). Likewise, cardiac myocyte cross-section area measured in wheat-germ agglutinin-stained LV tissue sections was normalized in TM54 mice expressing ONM-Nb80, when compared to the two control TM54 cohorts and wildtype littermates (Figure 7N,O). In addition, the interstitial myocardial fibrosis associated with DCM was inhibited by ONM-Nb80 treatment (Figure 7P,Q). These results indicate that treatment with an ONM-Nb80 gene therapy vector improved the cardiac structure and function of the TM54 mouse. Taken together, results obtained by expression of OMN-ICL3-9 and OMN-Nb80 *in vivo* demonstrate the essential role of perinuclear βARs in pathological cardiac remodeling, while providing proof-of-concept for the targeting of this GPCR compartment in dilated cardiomyopathy.

## Discussion

This study provides molecular and physiological evidence of an autonomous, nanometer scale, GPCR compartment important for determination of cellular structure and function. AKAP6 organizes perinuclear multimolecular signalosomes, which in response to locally generated and restricted cAMP signals, regulate pathological cardiac myocyte hypertrophy, retinal ganglion cell neuroprotection, and skeletal muscle regeneration.^16,49,50^ Here we define βARs located on Golgi membranes facing the ONM as uniquely responsible for activation of these signalosomes, as studied in the context of pathological cardiac myocyte hypertrophy. Using an ONM localized activator and inhibitor of the βAR (ONM-ICL3-9 and ONM-Nb80), as well as a peptide that can disrupt AKAP6-AKAP9-mediated tethering of the Golgi to the ONM (SR1-mCherry), we demonstrate that, in contrast to βAR on endosomes or the plasma membrane, Golgi-localized βAR stimulates perinuclear cAMP signaling, independently of cAMP signaling elsewhere in the cell. Perinuclear βARs thereby regulate perinuclear [Ca^2+^], activation of the calcineurin-NFAT pathway, myocyte hypertrophy, and ANF expression. The local regulation of AKAP6 signalosomes by perinuclear βARs permits the modulation of AKAP6-dependent signal transduction independently of other important βAR-dependent cellular processes in the myocyte, such as contractility, with biological relevance provided by perinuclear βAR gain- and loss-of-function in mouse models of cardiomyopathy. Taken together, this study demonstrates how compartmentalization provides specificity to GPCR signaling, illustrating how an understanding of the architecture of cellular compartmentation can be leveraged to alter selectivity cellular function, with potentially beneficial therapeutic effect.

There has been much recent interest in the functional significance of GPCRs detected within the cell away from the plasma membrane, in particular, the potential initiation of signal transduction at intracellular sites. In the cardiac myocyte, βARs have been detected on endosomes, Golgi apparatus, sarcoplasmic reticulum, and nuclear envelope, and the function of these internal receptors has been demonstrated using a pharmacological approach similar to that used in the initial experiments described herein.^5–8^ In contrast to endosomal GPCRs, Golgi-localized β_1_AR apparently signal without prior internalization from the plasma membrane.^7^ Golgi-resident β_1_AR activated by OCT3-transported NE has been shown to induce neonatal myocyte hypertrophy via Epac-mediated stimulation of phospholipase ε (PLCε), that can hydrolyze phosphatidylinositol-4-phosphate (PI4P) to diacylglycerol and inositol bisphosphate (IP_2_).^5^ PLCε can bind AKAP6β, regulating histone deacetylase nuclear export.^51^ Because of the inhibition of PI4P hydrolysis by a PLCε-derived peptide capable of binding AKAP6β and inhibiting PLCε-AKAP6β association, it was suggested that these Golgi-localized β_1_ARs activate PLCε via AKAP6β-bound Epac1.^5^ However, PI4P hydrolysis was reported to be insensitive to isoproterenol that stimulated cAMP production,^52^ while we have found that the OMN-AKAR4 sensor is robustly activated in myocytes by isoproterenol (at the same 1 µM dose).^27^ As PIP4 was assayed using a biosensor (FAPP-PH-GFP) that labels the Golgi broadly,^51^ it is possible that different pools of Golgi-associated β_1_AR and cAMP regulate AKAP6β signalosomes and Golgi PI4P hydrolysis. Like the plasma membrane that contains multiple discrete GPCR compartments,^53^ multiple separate GPCR compartments likely exist within the multilayered Golgi apparatus, regulating different signaling pathways, potentially under the control of different phosphodiesterases.^52^ Results obtained with ONM-Nb80 and ONM-ICL3-9, which can bridge the outer nuclear envelope and Golgi, suggest that βARs regulating the AKAP6β cAMP compartment are on a limited section of the Golgi directly facing the ONM. Further research will be required to define the role of AKAP6β in the regulation of PLCε by Golgi-resident βARs.

While targeting of the ICL3-9 pepducin to the plasma membrane induced PKA activity detected only with the cytosolic parental AKAR4 sensor, targeting of ICL3-9 to the Golgi and ONM induced PKA only at the nuclear envelope, thereby supporting the independence of the two signaling compartments. Interestingly, constitutive activation of surface βAR with PM-ICL3-9 also lowered steady-state ONM-AKAR signals (Figure 3E), an effect observed to a lesser degree with Endo-ICL3-9. How G_αs_-biased signaling by chronically activated surface βARs might down-regulate basal PKA signaling at perinuclear AKAP6β signalosomes is unclear, although potentially implicating plasma membrane βARs in the normal suppression of AKAP6β pathological signaling. Regardless, PM-ICL3-9 was unable to inhibit NE-induced ANF expression (Figure 6K). Instead, ANF was robustly regulated by ONM-Nb80 and ONM-ICL3-9, supporting further an independent, dominant function of the perinuclear compartment in myocyte hypertrophy.

A fundamental question in the field of signal transduction is how large are signaling compartments. Anton, et al, have shown that in HEK293 cells glucagon-like peptide 1 (GLP-1) receptors control “receptor-associated independent cAMP nanodomains” (RAINs) with a radius of ∼60 nm from the plasma membrane receptor.^53^ Images obtained by electron microscopy suggest that Golgi membrane can be within 100 nm of the ONM.^54^ It is likely that at AKAP6β signalosomes, the Golgi and ONM are even closer, as the ONM-Nb80 and ONM-ICL3-9 polypeptides were able to bridge the ONM and Golgi βARs. In addition, the N-terminal domain of AKAP6β binds directly adenylyl cyclase 5.^26^ G_α_ proteins, including G_αs_, have long been recognized to reside on the Golgi.^55^ Due to the activation of adenylyl cyclase by G_αs_, we suggest that the cyclase is located near βAR on the Golgi facing the nuclear envelope. As AKAP6 binds the conserved catalytic C1 and C2 domains of adenylyl cyclase 5,^26^ and Nb80 binds the cytoplasmic end of the βAR,^56^ both βAR and adenylyl cyclase 5 are presumably oriented with their cytoplasmic domains within the cleft between the Golgi and ONM. Thus, endogenous catecholamines would have to traverse both the plasma membrane and Golgi to bind AKAP6β signalosome-associated βARs within the Golgi lumen. NE presumably crosses both membranes via OCT3, which has been detected on both the plasma membrane and intracellular membranes.^7^ The remarkably close proximity of the ONM and Golgi at AKAP6β supports our hypothesis that AKAP6β organizes a highly insulated, independent signaling nano-compartment located between the Golgi and ONM.

Within the Golgi-ONM cleft, we detected a steep cAMP gradient by comparing signals obtained with ONM-Epac2-camps and ONM-50-Epac2-camps, such that <26 nm away from the N-terminus of nesprin-1α cAMP was decreased sufficiently to preclude sensor activation. We suggest that this gradient is maintained primarily by two mechanisms involving AKAP6β binding partners. First, PKA regulatory subunits, which are in molar excess of PKA catalytic subunits in most cells, can serve as a buffer for ambient cAMP.^2^ As shown by expression of the SuperAKAP-*IS*-mCherry fusion protein, the binding of PKA RII-subunits to AKAP6β limited cAMP to that compartment. Second, the recruitment of PKA-activated PDE4D3 by AKAP6β provides for the local degradation of cAMP, limiting its overall accumulation. Accordingly, displacement of PDE4D3 also was sufficient to dissipate the cAMP gradient and to permit the detection of ONM-ICL3-induced cAMP by soluble Epac2-camps sensor. Additional mechanisms might also contribute to the small physical dimensions of the signaling compartment. For example, Golgi membrane and ONM presumably provide a physical barrier to cAMP diffusion from other sites of the myocyte. The direct binding of adenylyl cyclase 5 to AKAP6β provides an explanation how, despite this steep gradient, local cAMP levels can be sufficient to activate AKAP6β-bound PKA.^26^ Interestingly, it has been suggested that in cells activated by endogenous ligands, cAMP levels are insufficient to promote the dissociation of PKA regulatory and catalytic subunits and, instead, induce PKA holoenzyme to undergo conformational shifts permitting the phosphorylation of adjacent substrates.^57^ The range at which PKA bound to an AKAP might phosphorylate a substrate has been predicted to be of the same scale (up to 25 nm) as the limit of the cAMP gradient detected in the current study.^57^ These results are intriguing as, like PDE4D3, AKAP6β-bound adenylyl cyclase 5 itself is a target for phosphorylation by AKAP6β-bound PKA.^26^ These results imply that relevant substrates for AKAP6β-bound PKA are similarly near the kinase.

In comparison to adult ventricular myocytes, cultured primary neonatal rat ventricular myocytes have a rudimentary transverse tubule and sarcoplasmic reticulum system and sparse myofibrils, with notable differences in Ca^2+^ handling and excitation-contraction coupling.^31,58^ The differences in ultrastructure between neonatal and adult myocytes are in many ways similar to the differences between adult myocytes in the normal and failing heart and may explain why studies in neonatal myocytes have often been informative regarding the regulation of pathological cardiac remodeling.^11,59^ A remarkable feature of the current study, as well as many of our earlier AKAP6β signalosome studies, is that results regarding signaling by βAR in the perinuclear compartment were similar whether obtained in neonatal or adult myocytes, while recapitulated in live mouse models of cardiovascular disease. This conservation of function presumably reflects the close perinuclear location of the Golgi in both neonatal and adult myocytes and the highly restricted localization of AKAP6β in these cells. Despite early reports suggesting the presence of AKAP6β throughout the sarcoplasmic reticulum,^60,61^ AKAP6β (mAKAPβ) is localized to the nuclear envelope in terminally differentiated myocytes by binding nesprin-1α.^13,22,25,35^

Elevated plasma norepinephrine levels (2-10 nM in heart failure compared to ∼1 nM in normals) are a common finding among patients in heart failure and prior to the development of β-blocker therapies were highly predictive of mortality.^62^ Although β_1_AR signaling is overall down-regulated in heart failure, recent studies have found that β_1_AR signaling is differentially regulated in heart failure depending upon the myocyte compartment, including a shift from the plasma membrane to internal compartments.^9,63^ We suggest that the internal signaling compartment defined by βAR association with AKAP6β signalosomes is a critical target of β-blocker therapy. Complementing *in vitro* studies in primary myocytes, we found that expression of the ONM-ICL3-9 activating protein in the mouse heart at low levels rapidly induced systolic dysfunction and pathological remodeling in the absence of other primary disease. Based upon the requirement for activation of the AKAP6β compartment for myocyte hypertrophy, we considered that, conversely, inhibition of AKAP6β-associated perinuclear βARs would be efficacious for the treatment of heart failure, an approach avoiding the common side effects of β-blockers such as low heart rate (bradycardia) and blood pressure (hypotension). To test this hypothesis, we treated a mouse model of familial, non-ischemic Dilated Cardiomyopathy. Non-ischemic Dilated Cardiomyopathy, of which 30-50% cases are genetic, incurs a 3-year mortality of 12-20% due to heart failure and ventricular arrythmia despite modern therapy.^64^ Importantly, with a prevalence as high as 1 in 250 in the adult population,^65^ Dilated Cardiomyopathy comprises a significant cause of cardiovascular mortality and is the most common indication for heart transplantation.^66^ Remarkably, treatment of the TM54 model for Dilated Cardiomyopathy with a ONM-Nb80 gene therapy vector improved cardiac function and decreased pathological remodeling. Further research will be required to determine the promise of gene therapy targeting perinuclear βARs and to understand the long-term consequences of inhibiting GPCRs dedicated to the regulation of cellular stress responses. In the meantime, this study provides evidence for the unique role of Golgi-localized βAR in regulating an independent perinuclear cAMP nanodomain. This nanodomain is responsible for the induction of myocyte gene transcription and hypertrophy associated with pathological cardiac remodeling, providing a target for the development of novel compartment-specific beta-blockers for the treatment of cardiac disease.

## Acknowledgement and funding

This work was supported by the National Institutes of Health (HL153835 and HL166547 to KDK-D and HL158052 to MSK) and American Heart Association (PRE34030209 to MT).

## Author contribution

Moriah Turcotte: study design, data collection, analysis and interpretation, writing of manuscript; Sofia M. Possidento: data acquisition; Anne-Maj Samuelsson, Jinliang Li, and Zhuyun Qin: data collection, analysis and interpretation of in vivo studies; Kimberly Dodge-Kafka and Michael Kapiloff: project supervision, conception and design, data collection, analysis and interpretation, writing and final approval of manuscript.

## Data availability statement

The data that support the findings of this study are available from the corresponding authors upon request.

## Supplemental information

**Document S1.** Figures S1-S6

**Video S1.** Related to Figure 8. ONM-ICL3-9 treated C57BL/6N mouse, EF = 38%.

**Video S2.** Related to Figure 8. ONM-Control treated C57BL/6N mouse, EF = 53%.

**Video S3.** Related to Figure 8. C57BL/6N control mouse, EF = 49%.

**Video S4.** Related to Figure 8. OMN-Nb80 treated NTG mouse, EF = 49%.

**Video S5.** Related to Figure 8. OMN-Nb80 treated TM54 mouse, EF = 39%.

**Video S6.** Related to Figure 8. OMN-Control treated NTG mouse, EF = 46%.

**Video S7.** Related to Figure 8. OMN-Control treated TM54 mouse, EF = 17%.

**Video S8.** Related to Figure 8. NTG control mouse, EF = 46%.

**Video S9.** Related to Figure 8. TM54 control mouse, EF = 24%.

## References

1. Weis, W.I., and Kobilka, B.K. (2018). The Molecular Basis of G Protein-Coupled Receptor Activation. Annu Rev Biochem 87, 897–919. 10.1146/annurev-biochem-060614-033910.

2. Zaccolo, M., Zerio, A., and Lobo, M.J. (2021). Subcellular Organization of the cAMP Signaling Pathway. Pharmacol Rev 73, 278–309. 10.1124/pharmrev.120.000086.

3. Bock, A., Irannejad, R., and Scott, J.D. (2024). cAMP signaling: a remarkably regional affair. Trends Biochem Sci 49, 305–317. 10.1016/j.tibs.2024.01.004.

4. Collins, K.B., and Scott, J.D. (2023). Phosphorylation, compartmentalization, and cardiac function. IUBMB Life 75, 353–369. 10.1002/iub.2677.

5. Nash, C.A., Wei, W., Irannejad, R., and Smrcka, A.V. (2019). Golgi localized beta1-adrenergic receptors stimulate Golgi PI4P hydrolysis by PLCepsilon to regulate cardiac hypertrophy. Elife 8. 10.7554/eLife.48167.

6. Wang, Y., Shi, Q., Li, M., Zhao, M., Reddy Gopireddy, R., Teoh, J.P., Xu, B., Zhu, C., Ireton, K.E., Srinivasan, S., et al. (2021). Intracellular beta1-Adrenergic Receptors and Organic Cation Transporter 3 Mediate Phospholamban Phosphorylation to Enhance Cardiac Contractility. Circ Res 128, 246–261. 10.1161/CIRCRESAHA.120.317452.

7. Irannejad, R., Pessino, V., Mika, D., Huang, B., Wedegaertner, P.B., Conti, M., and von Zastrow, M. (2017). Functional selectivity of GPCR-directed drug action through location bias. Nat Chem Biol 13, 799–806. 10.1038/nchembio.2389.

8. Tadevosyan, A., Vaniotis, G., Allen, B.G., Hebert, T.E., and Nattel, S. (2012). G protein-coupled receptor signalling in the cardiac nuclear membrane: evidence and possible roles in physiological and pathophysiological function. J Physiol 590, 1313–1330. 10.1113/jphysiol.2011.222794.

9. Wei, W., and Smrcka, A.V. (2022). Subcellular beta-Adrenergic Receptor Signaling in Cardiac Physiology and Disease. J Cardiovasc Pharmacol 80, 334–341. 10.1097/FJC.0000000000001324.

10. Grassi, G., Quarti-Trevano, F., and Esler, M.D. (2021). Sympathetic activation in congestive heart failure: an updated overview. Heart Fail Rev 26, 173–182. 10.1007/s10741-019-09901-2.

11. Nakamura, M., and Sadoshima, J. (2018). Mechanisms of physiological and pathological cardiac hypertrophy. Nat Rev Cardiol 15, 387–407. 10.1038/s41569-018-0007-y.

12. Heidenreich, P.A., Bozkurt, B., Aguilar, D., Allen, L.A., Byun, J.J., Colvin, M.M., Deswal, A., Drazner, M.H., Dunlay, S.M., Evers, L.R., et al. (2022). 2022 AHA/ACC/HFSA Guideline for the Management of Heart Failure: A Report of the American College of Cardiology/American Heart Association Joint Committee on Clinical Practice Guidelines. Circulation 145, e895–e1032. 10.1161/CIR.0000000000001063.

13. Pare, G.C., Easlick, J.L., Mislow, J.M., McNally, E.M., and Kapiloff, M.S. (2005). Nesprin-1alpha contributes to the targeting of mAKAP to the cardiac myocyte nuclear envelope. Exp Cell Res 303, 388–399. 10.1016/j.yexcr.2004.10.009.

14. Becker, R., Vergarajauregui, S., Billing, F., Sharkova, M., Lippolis, E., Mamchaoui, K., Ferrazzi, F., and Engel, F.B. (2021). Myogenin controls via AKAP6 non-centrosomal microtubule-organizing center formation at the nuclear envelope. Elife 10. 10.7554/eLife.65672.

15. Holt, I., Fuller, H.R., Lam, L.T., Sewry, C.A., Shirran, S.L., Zhang, Q., Shanahan, C.M., and Morris, G.E. (2019). Nesprin-1-alpha2 associates with kinesin at myotube outer nuclear membranes, but is restricted to neuromuscular junction nuclei in adult muscle. Sci Rep 9, 14202. 10.1038/s41598-019-50728-6.

16. Dodge-Kafka, K., Gildart, M., Tokarski, K., and Kapiloff, M.S. (2019). mAKAPbeta signalosomes - A nodal regulator of gene transcription associated with pathological cardiac remodeling. Cell Signal 63, 109357. 10.1016/j.cellsig.2019.109357.

17. Li, J., Tan, Y., Passariello, C.L., Martinez, E.C., Kritzer, M.D., Li, X., Li, X., Li, Y., Yu, Q., Ohgi, K., et al. (2020). Signalosome-Regulated Serum Response Factor Phosphorylation Determining Myocyte Growth in Width Versus Length as a Therapeutic Target for Heart Failure. Circulation 142, 2138–2154. 10.1161/CIRCULATIONAHA.119.044805.

18. Passariello, C.L., Gayanilo, M., Kritzer, M.D., Thakur, H., Cozacov, Z., Rusconi, F., Wieczorek, D., Sanders, M., Li, J., and Kapiloff, M.S. (2013). p90 ribosomal S6 kinase 3 contributes to cardiac insufficiency in alpha-tropomyosin Glu180Gly transgenic mice. Am J Physiol Heart Circ Physiol 305, H1010–1019. 10.1152/ajpheart.00237.2013.

19. Dodge-Kafka, K.L., Soughayer, J., Pare, G.C., Carlisle Michel, J.J., Langeberg, L.K., Kapiloff, M.S., and Scott, J.D. (2005). The protein kinase A anchoring protein mAKAP coordinates two integrated cAMP effector pathways. Nature 437, 574–578. 10.1038/nature03966.

20. Li, J., Kritzer, M.D., Michel, J.J., Le, A., Thakur, H., Gayanilo, M., Passariello, C.L., Negro, A., Danial, J.B., Oskouei, B., et al. (2013). Anchored p90 ribosomal S6 kinase 3 is required for cardiac myocyte hypertrophy. Circ Res 112, 128–139. 10.1161/CIRCRESAHA.112.276162.

21. Pare, G.C., Bauman, A.L., McHenry, M., Michel, J.J., Dodge-Kafka, K.L., and Kapiloff, M.S. (2005). The mAKAP complex participates in the induction of cardiac myocyte hypertrophy by adrenergic receptor signaling. J Cell Sci 118, 5637–5646. 10.1242/jcs.02675.

22. Kritzer, M.D., Li, J., Passariello, C.L., Gayanilo, M., Thakur, H., Dayan, J., Dodge-Kafka, K., and Kapiloff, M.S. (2014). The scaffold protein muscle A-kinase anchoring protein beta orchestrates cardiac myocyte hypertrophic signaling required for the development of heart failure. Circ Heart Fail 7, 663–672. 10.1161/CIRCHEARTFAILURE.114.001266.

23. Martinez, E.C., Li, J., Ataam, J.A., Tokarski, K., Thakur, H., Karakikes, I., Dodge-Kafka, K., and Kapiloff, M.S. (2023). Targeting mAKAPbeta expression as a therapeutic approach for ischemic cardiomyopathy. Gene Ther 30, 543–551. 10.1038/s41434-022-00321-w.

24. Dodge, K.L., Khouangsathiene, S., Kapiloff, M.S., Mouton, R., Hill, E.V., Houslay, M.D., Langeberg, L.K., and Scott, J.D. (2001). mAKAP assembles a protein kinase A/PDE4 phosphodiesterase cAMP signaling module. EMBO J 20, 1921–1930. 10.1093/emboj/20.8.1921.

25. Kapiloff, M.S., Schillace, R.V., Westphal, A.M., and Scott, J.D. (1999). mAKAP: an A-kinase anchoring protein targeted to the nuclear membrane of differentiated myocytes. J Cell Sci 112 ( *Pt 16*), 2725–2736. 10.1242/jcs.112.16.2725.

26. Kapiloff, M.S., Piggott, L.A., Sadana, R., Li, J., Heredia, L.A., Henson, E., Efendiev, R., and Dessauer, C.W. (2009). An adenylyl cyclase-mAKAPbeta signaling complex regulates cAMP levels in cardiac myocytes. J Biol Chem 284, 23540–23546. 10.1074/jbc.M109.030072.

27. Turcotte, M.G., Thakur, H., Kapiloff, M.S., and Dodge-Kafka, K.L. (2022). A perinuclear calcium compartment regulates cardiac myocyte hypertrophy. J Mol Cell Cardiol 172, 26–40. 10.1016/j.yjmcc.2022.07.007.

28. Boczek, T., Cameron, E.G., Yu, W., Xia, X., Shah, S.H., Castillo Chabeco, B., Galvao, J., Nahmou, M., Li, J., Thakur, H., et al. (2019). Regulation of Neuronal Survival and Axon Growth by a Perinuclear cAMP Compartment. J Neurosci 39, 5466–5480. 10.1523/JNEUROSCI.2752-18.2019.

29. Wright, C.D., Chen, Q., Baye, N.L., Huang, Y., Healy, C.L., Kasinathan, S., and O’Connell, T.D. (2008). Nuclear alpha1-adrenergic receptors signal activated ERK localization to caveolae in adult cardiac myocytes. Circ Res 103, 992–1000. 10.1161/CIRCRESAHA.108.176024.

30. Khanppnavar, B., Maier, J., Herborg, F., Gradisch, R., Lazzarin, E., Luethi, D., Yang, J.W., Qi, C., Holy, M., Jantsch, K., et al. (2022). Structural basis of organic cation transporter-3 inhibition. Nat Commun 13, 6714. 10.1038/s41467-022-34284-8.

31. Parra, V., and Rothermel, B.A. (2017). Calcineurin signaling in the heart: The importance of time and place. J Mol Cell Cardiol 103, 121–136. 10.1016/j.yjmcc.2016.12.006.

32. Staus, D.P., Strachan, R.T., Manglik, A., Pani, B., Kahsai, A.W., Kim, T.H., Wingler, L.M., Ahn, S., Chatterjee, A., Masoudi, A., et al. (2016). Allosteric nanobodies reveal the dynamic range and diverse mechanisms of G-protein-coupled receptor activation. Nature 535, 448–452. 10.1038/nature18636.

33. Staus, D.P., Wingler, L.M., Strachan, R.T., Rasmussen, S.G., Pardon, E., Ahn, S., Steyaert, J., Kobilka, B.K., and Lefkowitz, R.J. (2014). Regulation of beta2-adrenergic receptor function by conformationally selective single-domain intrabodies. Mol Pharmacol 85, 472–481. 10.1124/mol.113.089516.

34. Carr, R., 3rd, Du, Y., Quoyer, J., Panettieri, R.A., Jr., Janz, J.M., Bouvier, M., Kobilka, B.K., and Benovic, J.L. (2014). Development and characterization of pepducins as Gs-biased allosteric agonists. J Biol Chem 289, 35668–35684. 10.1074/jbc.M114.618819.

35. Vergarajauregui, S., Becker, R., Steffen, U., Sharkova, M., Esser, T., Petzold, J., Billing, F., Kapiloff, M.S., Schett, G., Thievessen, I., and Engel, F.B. (2020). AKAP6 orchestrates the nuclear envelope microtubule-organizing center by linking golgi and nucleus via AKAP9. Elife 9. 10.7554/eLife.61669.

36. Escobar, M., Cardenas, C., Colavita, K., Petrenko, N.B., and Franzini-Armstrong, C. (2011). Structural evidence for perinuclear calcium microdomains in cardiac myocytes. J Mol Cell Cardiol 50, 451–459. 10.1016/j.yjmcc.2010.11.021.

37. Gillooly, D.J., Morrow, I.C., Lindsay, M., Gould, R., Bryant, N.J., Gaullier, J.M., Parton, R.G., and Stenmark, H. (2000). Localization of phosphatidylinositol 3-phosphate in yeast and mammalian cells. EMBO J 19, 4577–4588. 10.1093/emboj/19.17.4577.

38. Fraser, I.D., Tavalin, S.J., Lester, L.B., Langeberg, L.K., Westphal, A.M., Dean, R.A., Marrion, N.V., and Scott, J.D. (1998). A novel lipid-anchored A-kinase Anchoring Protein facilitates cAMP-responsive membrane events. EMBO J 17, 2261–2272. 10.1093/emboj/17.8.2261.

39. Wu, X., and Bers, D.M. (2006). Sarcoplasmic reticulum and nuclear envelope are one highly interconnected Ca2+ store throughout cardiac myocyte. Circ Res 99, 283–291. 10.1161/01.RES.0000233386.02708.72.

40. Nikolaev, V.O., Bunemann, M., Hein, L., Hannawacker, A., and Lohse, M.J. (2004). Novel single chain cAMP sensors for receptor-induced signal propagation. J Biol Chem 279, 37215–37218. 10.1074/jbc.C400302200.

41. Arai, R., Ueda, H., Kitayama, A., Kamiya, N., and Nagamune, T. (2001). Design of the linkers which effectively separate domains of a bifunctional fusion protein. Protein Eng 14, 529–532.

42. Gold, M.G., Lygren, B., Dokurno, P., Hoshi, N., McConnachie, G., Tasken, K., Carlson, C.R., Scott, J.D., and Barford, D. (2006). Molecular basis of AKAP specificity for PKA regulatory subunits. Mol Cell 24, 383–395. 10.1016/j.molcel.2006.09.006.

43. Chen, T.W., Wardill, T.J., Sun, Y., Pulver, S.R., Renninger, S.L., Baohan, A., Schreiter, E.R., Kerr, R.A., Orger, M.B., Jayaraman, V., et al. (2013). Ultrasensitive fluorescent proteins for imaging neuronal activity. Nature 499, 295–300. 10.1038/nature12354.

44. Li, J., Negro, A., Lopez, J., Bauman, A.L., Henson, E., Dodge-Kafka, K., and Kapiloff, M.S. (2010). The mAKAPbeta scaffold regulates cardiac myocyte hypertrophy via recruitment of activated calcineurin. J Mol Cell Cardiol 48, 387–394. 10.1016/j.yjmcc.2009.10.023.

45. Mehta, S., Aye-Han, N.N., Ganesan, A., Oldach, L., Gorshkov, K., and Zhang, J. (2014). Calmodulin-controlled spatial decoding of oscillatory Ca2+ signals by calcineurin. Elife 3, e03765. 10.7554/eLife.03765.

46. Sergeeva, I.A., and Christoffels, V.M. (2013). Regulation of expression of atrial and brain natriuretic peptide, biomarkers for heart development and disease. Biochim Biophys Acta 1832, 2403–2413. 10.1016/j.bbadis.2013.07.003.

47. Prasad, K.M., Xu, Y., Yang, Z., Acton, S.T., and French, B.A. (2011). Robust cardiomyocyte-specific gene expression following systemic injection of AAV: in vivo gene delivery follows a Poisson distribution. Gene Ther 18, 43–52. 10.1038/gt.2010.105.

48. Rajan, S., Ahmed, R.P., Jagatheesan, G., Petrashevskaya, N., Boivin, G.P., Urboniene, D., Arteaga, G.M., Wolska, B.M., Solaro, R.J., Liggett, S.B., and Wieczorek, D.F. (2007). Dilated cardiomyopathy mutant tropomyosin mice develop cardiac dysfunction with significantly decreased fractional shortening and myofilament calcium sensitivity. Circ Res 101, 205–214. 10.1161/CIRCRESAHA.107.148379.

49. Boczek, T., Yu, Q., Zhu, Y., Dodge-Kafka, K.L., Goldberg, J.L., and Kapiloff, M.S. (2021). cAMP at Perinuclear mAKAPalpha Signalosomes Is Regulated by Local Ca(2+) Signaling in Primary Hippocampal Neurons. eNeuro 8. 10.1523/ENEURO.0298-20.2021.

50. Lee, S.W., Won, J.Y., Yang, J., Lee, J., Kim, S.Y., Lee, E.J., and Kim, H.S. (2015). AKAP6 inhibition impairs myoblast differentiation and muscle regeneration: Positive loop between AKAP6 and myogenin. Sci Rep 5, 16523. 10.1038/srep16523.

51. Zhang, L., Malik, S., Pang, J., Wang, H., Park, K.M., Yule, D.I., Blaxall, B.C., and Smrcka, A.V. (2013). Phospholipase Cepsilon hydrolyzes perinuclear phosphatidylinositol 4-phosphate to regulate cardiac hypertrophy. Cell 153, 216–227. 10.1016/j.cell.2013.02.047.

52. Nash, C.A., Brown, L.M., Malik, S., Cheng, X., and Smrcka, A.V. (2018). Compartmentalized cyclic nucleotides have opposing effects on regulation of hypertrophic phospholipase Cepsilon signaling in cardiac myocytes. J Mol Cell Cardiol 121, 51–59. 10.1016/j.yjmcc.2018.06.002.

53. Anton, S.E., Kayser, C., Maiellaro, I., Nemec, K., Moller, J., Koschinski, A., Zaccolo, M., Annibale, P., Falcke, M., Lohse, M.J., and Bock, A. (2022). Receptor-associated independent cAMP nanodomains mediate spatiotemporal specificity of GPCR signaling. Cell 185, 1130–1142 e1111. 10.1016/j.cell.2022.02.011.

54. Yang, Z., Kirton, H.M., MacDougall, D.A., Boyle, J.P., Deuchars, J., Frater, B., Ponnambalam, S., Hardy, M.E., White, E., Calaghan, S.C., et al. (2015). The Golgi apparatus is a functionally distinct Ca2+ store regulated by the PKA and Epac branches of the beta1-adrenergic signaling pathway. Sci Signal 8, ra101. 10.1126/scisignal.aaa7677.

55. Denker, S.P., McCaffery, J.M., Palade, G.E., Insel, P.A., and Farquhar, M.G. (1996). Differential distribution of alpha subunits and beta gamma subunits of heterotrimeric G proteins on Golgi membranes of the exocrine pancreas. J Cell Biol 133, 1027–1040. 10.1083/jcb.133.5.1027.

56. Rasmussen, S.G., Choi, H.J., Fung, J.J., Pardon, E., Casarosa, P., Chae, P.S., Devree, B.T., Rosenbaum, D.M., Thian, F.S., Kobilka, T.S., et al. (2011). Structure of a nanobody-stabilized active state of the beta(2) adrenoceptor. Nature 469, 175–180. 10.1038/nature09648.

57. Smith, F.D., Esseltine, J.L., Nygren, P.J., Veesler, D., Byrne, D.P., Vonderach, M., Strashnov, I., Eyers, C.E., Eyers, P.A., Langeberg, L.K., and Scott, J.D. (2017). Local protein kinase A action proceeds through intact holoenzymes. Science 356, 1288–1293. 10.1126/science.aaj1669.

58. Poindexter, B.J., Smith, J.R., Buja, L.M., and Bick, R.J. (2001). Calcium signaling mechanisms in dedifferentiated cardiac myocytes: comparison with neonatal and adult cardiomyocytes. Cell Calcium 30, 373–382. 10.1054/ceca.2001.0249.

59. Louch, W.E., Koivumaki, J.T., and Tavi, P. (2015). Calcium signalling in developing cardiomyocytes: implications for model systems and disease. J Physiol 593, 1047–1063. 10.1113/jphysiol.2014.274712.

60. Yang, J., Drazba, J.A., Ferguson, D.G., and Bond, M. (1998). A-kinase anchoring protein 100 (AKAP100) is localized in multiple subcellular compartments in the adult rat heart. J Cell Biol 142, 511–522.

61. Marx, S.O., Reiken, S., Hisamatsu, Y., Jayaraman, T., Burkhoff, D., Rosemblit, N., and Marks, A.R. (2000). PKA phosphorylation dissociates FKBP12.6 from the calcium release channel (ryanodine receptor): defective regulation in failing hearts. Cell 101, 365–376. 10.1016/s0092-8674(00)80847-8.

62. Cohn, J.N., Levine, T.B., Olivari, M.T., Garberg, V., Lura, D., Francis, G.S., Simon, A.B., and Rector, T. (1984). Plasma norepinephrine as a guide to prognosis in patients with chronic congestive heart failure. N Engl J Med 311, 819–823. 10.1056/NEJM198409273111303.

63. Xu, B., Bahriz, S., Salemme, V.R., Wang, Y., Zhu, C., Zhao, M., and Xiang, Y.K. (2024). Differential Downregulation of beta(1)-Adrenergic Receptor Signaling in the Heart. J Am Heart Assoc 13, e033733. 10.1161/JAHA.123.033733.

64. Halliday, B.P., Cleland, J.G.F., Goldberger, J.J., and Prasad, S.K. (2017). Personalizing Risk Stratification for Sudden Death in Dilated Cardiomyopathy: The Past, Present, and Future. Circulation 136, 215–231. 10.1161/CIRCULATIONAHA.116.027134.

65. McKenna, W.J., and Judge, D.P. (2021). Epidemiology of the inherited cardiomyopathies. Nat Rev Cardiol 18, 22–36. 10.1038/s41569-020-0428-2.

66. Repetti, G.G., Toepfer, C.N., Seidman, J.G., and Seidman, C.E. (2019). Novel Therapies for Prevention and Early Treatment of Cardiomyopathies. Circ Res 124, 1536–1550. 10.1161/CIRCRESAHA.119.313569.

67. Depry, C., Allen, M.D., and Zhang, J. (2011). Visualization of PKA activity in plasma membrane microdomains. Mol Biosyst 7, 52–58. 10.1039/c0mb00079e.

68. Duong, N.T., Morris, G.E., Lam le, T., Zhang, Q., Sewry, C.A., Shanahan, C.M., and Holt, I. (2014). Nesprins: tissue-specific expression of epsilon and other short isoforms. PLoS One 9, e94380. 10.1371/journal.pone.0094380.

69. Randles, K.N., Lam le, T., Sewry, C.A., Puckelwartz, M., Furling, D., Wehnert, M., McNally, E.M., and Morris, G.E. (2010). Nesprins, but not sun proteins, switch isoforms at the nuclear envelope during muscle development. Dev Dyn 239, 998–1009. 10.1002/dvdy.22229.

70. Ring, A.M., Manglik, A., Kruse, A.C., Enos, M.D., Weis, W.I., Garcia, K.C., and Kobilka, B.K. (2013). Adrenaline-activated structure of beta2-adrenoceptor stabilized by an engineered nanobody. Nature 502, 575–579. 10.1038/nature12572.

