## Supplemental for "The Unique Role of Intracellular Perinuclear β-Adrenergic Receptors in defining Signaling Compartmentation and Pathological Cardiac Remodeling"

### Materials and methods

#### Animal Studies

All research in this manuscript was approved by the Institutional Animal Care and Use Committee at the University of Connecticut Health Center and conformed to the NIH Guide for the Care and Use of Laboratory Animals. Mouse research was approved by the Administrative Panel on Laboratory Animal Care (APLAC) Institutional Animal Care and Use Committee (IACUC) at Stanford University. Neonatal and adult rat cardiac myocytes were isolated from Sprague Dawley Rats purchased from Charles River. FVB/N-Tg(Myh6-Tpm1*E54K) "TM54" mice were previously provided by Dr. Beata Wolska and Dr. David Wieczorek and are available as strain #035610 at the Jackson Laboratory (Bar Harbor, ME).[1] Mice were genotyped by polymerase chain reaction using the following primers: Myh6 Forward: 5'-GCC CAC ACC AGA AAT GAC AGA-3' and Tpm1 Reverse: 5'-TCC AGT TCA TCT TCA GTG CCC-3' (236 bp product); *Atp1a2* internal control primers sense: 5'-AGC GAG CTC AGG ACA TTC TGG-3' and antisense: 5'-CTC CTA ACC ACG CTC CTA GCA-3' (494 bp).

#### Plasmids

All plasmids were constructed by Azenta Life Sciences (Genewiz, South Plainfield, NJ) or Vectorbuilder (Chicago, IL) using the methods of their choice. All plasmids were validated by sequencing, with most completely sequenced by next-generation sequencing, and by restriction digest before use. Complete plasmid sequences are available upon request. EGFP-GalT was a gift from Jennifer Lippincott-Schwartz (Addgene plasmid # 11929; http://n2t.net/addgene:11929 ; RRID:Addgene_11929).[2]

pS-AKAR4-nesprin and pS-AKAR4 encoding ONM-AKAR4 and parent AKAR4 were as previously described.[3] All pS series vectors direct cDNA expression under control of the cytomegalovirus immediate early promoter and contain I-Ceu I and PI-Sce I flanking sites for subcloning in the Adeno-X vector (Clontech Adeno-X Tet-Off Expression System 1). pTRE-GcAMP6S (containing the conditional TRE promoter and requiring co-infection with Adeno-tTA virus, Clontech Adeno-X Tet-Off Expression System 1), pS-GCaMP6s-nesprin, pS-CaNAR2, and pS-CaNAR2-nesprin were as previously described.[4] ONM-Epac2-camps was expressed using pS-Epac2-camps-nesprin that was identical to pS-CaNAR2-nesprin and pS-GCaMP6s-nesprin except for the substitution of a Flag-tagged Epac2-camps open reading frame from pCDNA3-epac2-camps (gift of Dr. Viacheslav Nikolaev).[5] ONM-50-Epac2-camps was expressed using pS-Epac2-camps-EAAAK50-nesprin-1 that contained a BssH II – Not I cassette encoding 50 EAAAK repeats between the Epac2-camps and nesprin-1 cDNAs. These sensor plasmids contain a myc-tagged human nesprin-1 cDNA (NCBI AF495910 bp 23997-26993), which is identical to the mRNA for nesprin-1α2 (the major isoform in cardiac myocytes, NCBI AY184203) except for substitution of a 76 aa sequence from the longer nesprin-1 isoforms in lieu of the first 31 aa of 1α2 and for an exclusion of the DV23 alternatively spliced 23 aa exon (of unknown function) generally present in heart.[6,7]

ONM-ICL3-9 and ONM-Nb80 were expressed using plasmids derived from the previously described control pS-mCherry-nesprin, in which a mCherry cDNA is 5' to the same nesprin-1 cDNA in the biosensor plasmids.[3] For ONM-ICL3-9, a cDNA fragment encoding a myc tag, the ICL3-9 peptide GRFHVQNLSQVEQDGRTIGII,[8] and a flexible $(GGGGS)_{13}$ linker was subcloned 5' to the mCherry cDNA sequence. For ONM-Nb80, a cDNA fragment encoding a myc tag, the Nb80 nanobody,[9] and a flexible $(GGGGS)_{13}$ linker was subcloned 5' to the mCherry cDNA sequence. PM-ICL3-9 was expressed using pS-AKAP18(1-25)-ICL3-9-mCherry-Flag in which a cDNA encodes (1) the first 25 aa of AKAP18α (MGQLCCFPFSRDEGKISEKNGGEPD),[10] (2) a flexible $(GGGGS)_{13}$ linker, (3) the ICL3-9 peptide, (4) mCherry, and (5) a C-terminal Flag tag. Endo-ICL3-9 was expressed using pS-FYVE-ICL3-9-mCherry-Flag in which a cDNA encodes (1) two direct repeats of mouse HGF-regulated tyrosine kinase substrate aa 147-223 (NCBI AAH03239) separated by QGQGS, (2) a flexible $(GGGGS)_{13}$ linker, (3) the ICL3-9 peptide, (4) mCherry, and (5) a C-terminal Flag tag. Golgi-ICL3-9 was expressed using pS-ICL3-9-GalT-mCherry-Flag in which a cDNA encodes (1) the ICL3-9 peptide, (2) a flexible $(GGGGS)_{13}$ linker, (3) Homo sapiens beta-1,4-galactosyltransferase 1 aa 2-82 (NCBI CDJ98633),[11] (4) mCherry, and (5) a C-terminal Flag tag. PM-ICL3-9, Endo-Nb80, and Golgi-Nb80 were expressed with the similar vectors pS-AKAP18(1-25)-Nb80-mCherry-Flag, pS-FYVE-Nb80-mCherry-Flag, and pS-Nb80-GalT-mCherry-Flag, respectively, in which the ICL3-9 peptide sequence was replaced with a Nb80 cDNA. AKAPIS-mCherry was expressed using pS-SuperAKAPIS-mCherry-Flag in which a cDNA encodes (1) the PKA binding peptide QIEYVAKQIVDYAIHQA,[12] (2) a flexible $(GGGGS)_{13}$ linker, (3) mCherry, and (5) a C-terminal Flag tag. 4D3(E)-mCherry was expressed using pscS2-4D3(E)-mCherry-mh, as previously described.[3] SR1-mCherry was expressed using a pS vector containing a cDNA for mAKAP aa 586-915 fused to Flag-tagged mCherry.

AAV shuttle plasmids were constructed with pAcTnTs (gift from Dr. Brent French) that directs expression under the control of the 407 bp chicken cardiac troponin T promoter.[13] OMN-Control AAV were generated using pAcTnT-myc-nesprin-1 that expresses the aforementioned myc-tagged nesprin-1 cDNA. OMN-Nb80 AAV were generated using pAcTnT-Nb80-myc-nesprin-1 that encodes Nb80 and a flexible $(GGGGS)_{13}$ linker N-terminal to myc-tagged nesprin-1.

Similarly, OMN-ICL3-9 AAV were generated using pAcTnT-ICL3-9-myc-nesprin-1 that encodes the ICL3-9 peptide and a flexible $(GGGGS)_{13}$ linker N-terminal to myc-tagged nesprin-1.

**Adenoviruses**

NFATc1-GFP was expressed using adenovirus obtained from Seven Hills Bioreagents (Catalog no. JMAd-98). All adenoviruses were generated by transfection of HEK293 cells with Adeno-X plasmids (Clontech Adeno-X Tet-Off Expression System 1) into which genes of interest were subcloned using the I-Ceu I and PI-Sce I restriction sites. Adenovirus were purified using Vivapure® AdenoPACK™ 20 kits (Sartorius), and titers were determined by end-point dilution method for HEK293 cell viability.

**Antibodies and Immunohistochemical Reagents**

| | | |
|---|---|---|
| α-actinin | mouse | Sigma monoclonal EA-53 |
| Atrial Natriuretic Peptide | rabbit | Sigma AB5490 |
| Atrial Natriuretic Peptide | Rabbit | Abcam ab225844, |
| Nesprin-1 | Rabbit | OR009 (custom)[14] |
| N-cadherin | Mouse | Thermofisher MA1-91128 |
| Alexa Fluor 555 wheat germ agglutinin conjugate | | Invitrogen W32464 |
| anti-mouse IgG (H+L) Alex Fluor 488 | goat | Thermofisher A-11001 |
| anti-rabbit IgG (H+L) Alex Fluor 488 | goat | Thermofisher A32731 |
| anti-rabbit IgG (H+L) Alexa Fluor 555 | donkey | Thermofisher A31572 |
| SlowFade Diamond Antifade Mountant with DAPI | | Thermofisher S36964 |
| Picrosirius Red Stain Kit | | Polysciences 24901-500 |

**Neonatal rat ventricular myocyte isolation and culture**

2-3-day-old Sprague-Dawley rats were euthanized by decapitation. Heart tissue was gradually dissociated through several rounds of collagenase wash and trituration followed by serum neutralization. Heart cells were collected by centrifugation and strained via 70 μm mesh cell strainer. Next, a 2-hour pre-plating period maximally removed fibroblasts, and the remaining myocytes were collected by centrifugation and plated on 1% gelatin-coated plates (500,000 myocytes per 35 mm plate) in Dulbecco's Modified Eagle Medium: Nutrient Mixture F-12 (DMEM/F12) supplemented with 1% penicillin/streptomycin (Gibco-BRL), 10% horse serum (HS), and 5% fetal bovine serum (FBS). The following day, cells were washed and cultured in serum-free DMEM/F12 containing antibiotic if being treated with adenovirus or DMEM/F12 containing

5% FBS if undergoing transfection. If applicable, 35 mm plates of RNV were transfected with Lipofectamine, 2 µg of each plasmid DNA and if needed, adenovirus co-infection the following day.

**Adult rat ventricular myocyte isolation and culture**

2-3-month-old male and female Sprague-Dawley rats were anti-coagulated by 300 U heparin intraperitoneal injection. 20-30 minutes later rats were anesthetized with ketamine (80 mg/kg) and xylazine (8 mg/kg) for heart excision. Hearts were collected into chilled perfusion buffer (mmol/L: NaCl 120, KCl 5.4, $Na_2HPO_4$ 1.2, $NaHCO_3$ 20, $MgCl_2$ 1.6, Taurine 5, Glucose 5.6, 2,3-Butanedione monoxime 10), equilibrated with 95% $O_2$ and 5% $CO_2$). The aorta was cannulated and perfused using a Harvard Langendorff apparatus with buffer at 37°C at a constant rate of 2.2 mL/min for 5 minutes, followed by perfusion for 45 minutes with 50 mL digestion buffer (perfusion buffer with 120 mg type II collagenase (Worthington, 315 U/mg), 5 mg protease (Sigma type XIV), and 55 mg BSA), recycling enzyme once reduced to 30 mL remaining. Perfusion was terminated when the hearts were soft enough for trituration. Atria were removed, and the ventricles cut into pieces before suspension and trituration in 5 mL digestion buffer with a large bore pipette, followed by filtration using 150-200µm nylon mesh. Cells were collected via centrifugation and subjected to gradual $Ca^{2+}$ stepwise addition (0.25, 0.5, and 1 mmol/L $Ca^{2+}$). The remaining myocytes were plated (100,000 myocytes per 35 mm plate) on laminin (Corning, 10 µg laminin per dish) coated plates followed by washing 1.5 hour later and addition of ACCT medium [Medium 199, 5 mmol/L Creatine (Sigma), 2 mmol/L L-carnitine (Sigma), 5 mmol/L Taurine (Sigma), 25 mmol/L Hepes (Sigma), 10 mmol/L 2, 3-Butanedione monoxime (ACROS Organics), 1% penicillin/streptomycin, 0.2% BSA (fatty acid free, Sigma), 0.1% ITS]. Adult myocytes were infected with adenovirus the same day as preparation, with drug treatments occurring the next day.

**Live Cell Imaging**

For imaging of both parent and ONM-targeted Epac2-camps, CaNAR2 and AKAR4 Forster resonance energy transfer (FRET) sensors and GCaMP6s intensiometric sensors, adult and neonatal myocytes were infected with adenovirus (multiplicity of infection [MOI] 5-50). Cells were imaged within two days after infection. For imaging, cells are washed and imaged in Hanks' Balanced Salt Solution (Gibco, mmol/L: $KH_2PO_4$ 0.44; $Na_2HPO_4$ 0.34; $NaHCO_3$ 4.2; NaCl 138, KCl 5.3; D-glucose 5.6; $CaCl_2$ 1.3; $MgCl_2$ 0.49; $MgSO_4$ 0.41). All imaging was done on a Zeiss Pascal confocal microscope using a 40x/1.2 numerical aperture objective, a 440 nm laser (Toptica

Photonics), and HQ535/50M and HQ480/40M emission and 510DCLP dichroic filters (Chroma Technology). For AKAR4 studies involving acute treatment with NE and serial imaging, images were acquired at 15 second intervals. For other sensors, images were acquired every 5 seconds. Due to a minimum required ~25 minutes for CaN to become active, cells expressing CaNAR2 or CaNAR2-nesprin were treated with drugs as indicated for at least 30 minutes prior to imaging. FRET for regions of interest was quantified using background-subtracted images and Image J, with statistical analyses performed using Graphpad Prism 8. FRET ratio "R" was defined as net FRET ÷ donor signal and normalized to $R_0$ (ratio for time=0) for experiments involving acute treatment. For Epac2-camps, which has decreased FRET signal upon cAMP binding,[5] 1/R was reported. For intensiometric GCaMP6s and GcaMP6s-nesprin sensors, fluorescent signal reflecting $Ca^{2+}$ binding was calculated for regions of interest using background-subtracted images and Image J with normalization to the fluorescence intensity of control samples. For each experiment, traces or single time point images were obtained for cells obtained from at least 3 different myocyte preparations.

The following drugs were used in this study: Norepinephrine (10 nM), Sotalol (20 μM), Propranolol (1 μM), Corticosterone (20 μM), H-89 (1 μM). All pharmacological interventions were added directly to the media of cells before initiation of live cell imaging, with corticosterone administered at least 20 minutes prior to imaging.

**Immunocytochemistry and cell based assays**

Myocytes were fixed using 3.7% formaldehyde in PBS for 10 minutes for neonatal and 1 hour for adult myocytes. Cells were permeabilized with 0.3% Triton X-100 in PBS and then washed and blocked using PBS containing 0.2% BSA and 1% horse serum for 30 minutes. Slides were incubated for 1 hour with primary antibodies, followed by 1-hour incubation with Alexa fluorescent dye-conjugated specific-secondary antibodies. Blocking buffer was used for antibody dilution and washes following primary and secondary antibody additions. Slides were mounted with SlowFade Diamond Mountant with DAPI. Widefield fluorescent images were acquired by a Zeiss Observer Z1 fluorescent microscope with an Axiocam camera.

NFATc1-GFP Localization and ANF expression assays: Neonatal myocytes were infected and/or transfected the day after plating as indicated. Cells were washed with maintenance medium and drugs added the following day with endpoint at 48 hours. For each slide, at least 50 cells were examined for NFATc1-GFP localization or 100 for perinuclear ANF staining. The relative localization of NFATc1-GFP in myocytes was measured as the ratio of nuclear vs cytosolic fluorescence intensity.

Myocyte Hypertrophy Assay: Adult myocytes were infected with adenovirus as indicated for 24 hours (MOI 5-50), followed by washing with ACCT media and treatment with drug as indicated for 48 hours. For each slide, at least 100 cells were measured for maximum length and width.

**Adeno-associated virus**

Serotype 9 AAV were produced by the University of Pennsylvania Vector Core and titered by ddPCR. AAV9 was injected via the tail vein ($5x10^{11}$ vg i.p.) into 6-8 week old male C57BL/6NJ mice (The Jackson Laboratory Strain #005304) or male and female TM54 and wildtype littermate mice as indicated. Masking of cohorts was provided by assigning a random number to each mouse by ear tag, such that identification of mouse cohort was not revealed until *in vivo* and post-mortem analyses were complete. A formal power analysis was not performed for the studies in this project. Mice were not selected for AAV delivery or expression before analysis.

**Echocardiography**

Mice, minimally anesthetized with 1-3% isoflurane, were studied by transthoracic echocardiography using a Vevo 3100 High-Resolution Imaging System (VisualSonics, Toronto, ON, Canada). A Visualsonic MX400 (20–46 MHz, 50 µm axial resolution) linear array transducer was used for all image acquisitions. For M-mode echocardiography, calculated parameters from at least three cardiac cycles were as follows: FS, fractional shortening = (LVID;d – LVID;s)/(LVID;d) in which LVID, LVAW, and LVPW are left ventricular interior diameter, anterior wall thickness and posterior wall thickness, respectively, and d and s refer to diastole and systole, respectively. For 4D image acquisition, the step motor was positioned just below the apex and the motor aligned to acquire concentric short axis images in 0.2 mm steps. At each position, a complete cardiac cycle was recorded using automated ECG and respiratory gating. 4D images were constructed using Vevo 4D image software. Image analysis was performed by blinded sonographer. 4D measurement of ejection fraction (EF) were calculated directly from volumetric measurements for end-diastolic (EDV) and end-systolic volume (ESV) based on operator-defined edge-tracing using Vevo 4D imaging software (VevoLAB, VisualSonics).

**Histochemistry**

Heart tissue was fixed in 3.7% formaldehyde. De-paraffinized 6 $\mu$m tissue sections were stained using the Picrosirius Red Stain Kit (Polysciences, Warrington, PA) and Alexa Fluor 555 wheat germ agglutinin conjugate (Invitrogen, Waltham, MA), as previously described.[15] For each mouse, the cross-section area of a total of >150 myocytes in >3 distinct regions of the left ventricle were measured using the wheat germ agglutinin-stained sections following fluorescent imaging at 200x using a Leica DM4000 microscope and a DFC3000G camera. Collagen content in myocardium, excluding the area around blood vessels, was assayed using Picrosirius Red stained sections and imaging of the entire left ventricle by circularly polarized light microscopy at 200x magnification using a Leica DM4000 microscope and a DMC2900 color camera, with illumination and exposure time settings optimized to differentiate bright collagen birefringence and the dark background. Measurements were acquired using NIH Image J or Leica LAS software.

**Statistical analysis**

Statistics were computed using Prism 10 (Graphpad, San Diego, California). All data are expressed as mean ± s.e.m. Unpaired, two-tailed t-tests and one-way or two-way ANOVA (with or without matching) followed by Tukey's (comparison with more than 2 groups for 1-way ANOVA or more than 2 groups in a row for 2-way ANOVA), Dunnett's (comparison of multiple groups with common control for 1-way ANOVA), or Uncorrected Fisher's LSD (comparisons of cell means with others in its row and its column for 2x2 design) post-hoc testing were performed as appropriate. Brown-Forsythe test was used to determine if there was a significant difference in standard deviation among multiple groups for 1-way ANOVA (as apparent by visual inspection for many of the imaging studies), and, if so, Brown-Forsythe and Welch ANOVA tests were used for ANOVA, with Dunnett's T3 test used for post-hoc testing. Likewise, Welch's correction was used for t-tests with significant differences in variances (F-test). Repeated symbols used as follows: * $p \leq 0.05$; ** $p \leq 0.01$; *** $p \leq 0.001$; **** $p \leq 0.0001$; ns - $p > 0.05$. $n$ refers to the number of individual mice, myocyte preparations, or live cell traces.

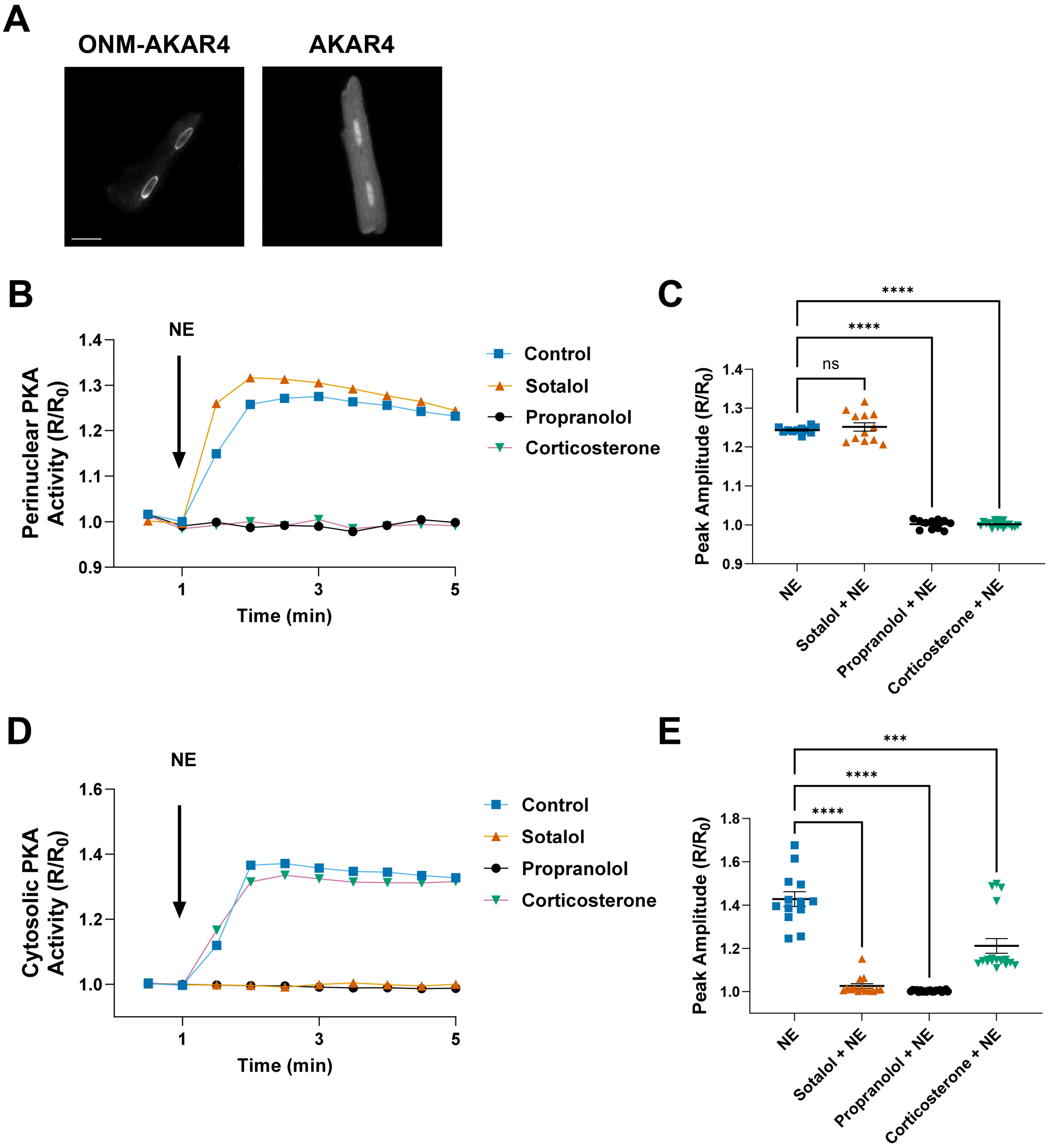

A
ONM-AKAR4
AKAR4
B
NE
Perinuclear PKA Activity (R/R0)
1.4
1.3
1.2
1.1
1.0
0.9
1
3
5
Time (min)
Control
Sotalol
Propranolol
Corticosterone
C
****
****
ns
Peak Amplitude (R/R0)
1.4
1.3
1.2
1.1
1.0
0.9
NE
Sotalol + NE
Propranolol + NE
Corticosterone + NE
D
NE
Cytosolic PKA Activity (R/R0)
1.6
1.4
1.2
1.0
1
3
5
Time (min)
Control
Sotalol
Propranolol
Corticosterone
E
***
****
****
Peak Amplitude (R/R0)
1.8
1.6
1.4
1.2
1.0
NE
Sotalol + NE
Propranolol + NE
Corticosterone + NE

**Supplemental Figure 1: An intracellular pool of β-adrenergic receptors activates AKAP6β-bound PKA in adult rat cardiomyocytes.**

(A) Adult rat ventricular myocytes (ARVM) were infected with either AKAR4 or ONM-AKAR4 adenovirus as indicated and imaged with a confocal microscope. Bar - 20 $\mu$m. Cyan image is shown in grayscale.

(B-E) FRET imaging of norepinephrine (NE)-treated ARVM expressing AKAR4 (B-C) or ONM-AKAR4 (D-E) and treated with either cell-impermeable β-blocker sotalol, cell-permeable β-blocker propranolol, or organic cation transporter 3 (Oct3) blocker corticosterone as indicated. Representative tracings and peak amplitudes for FRET ratio (R normalized to baseline $R_0$) are shown. P-values by Dunnett's test.

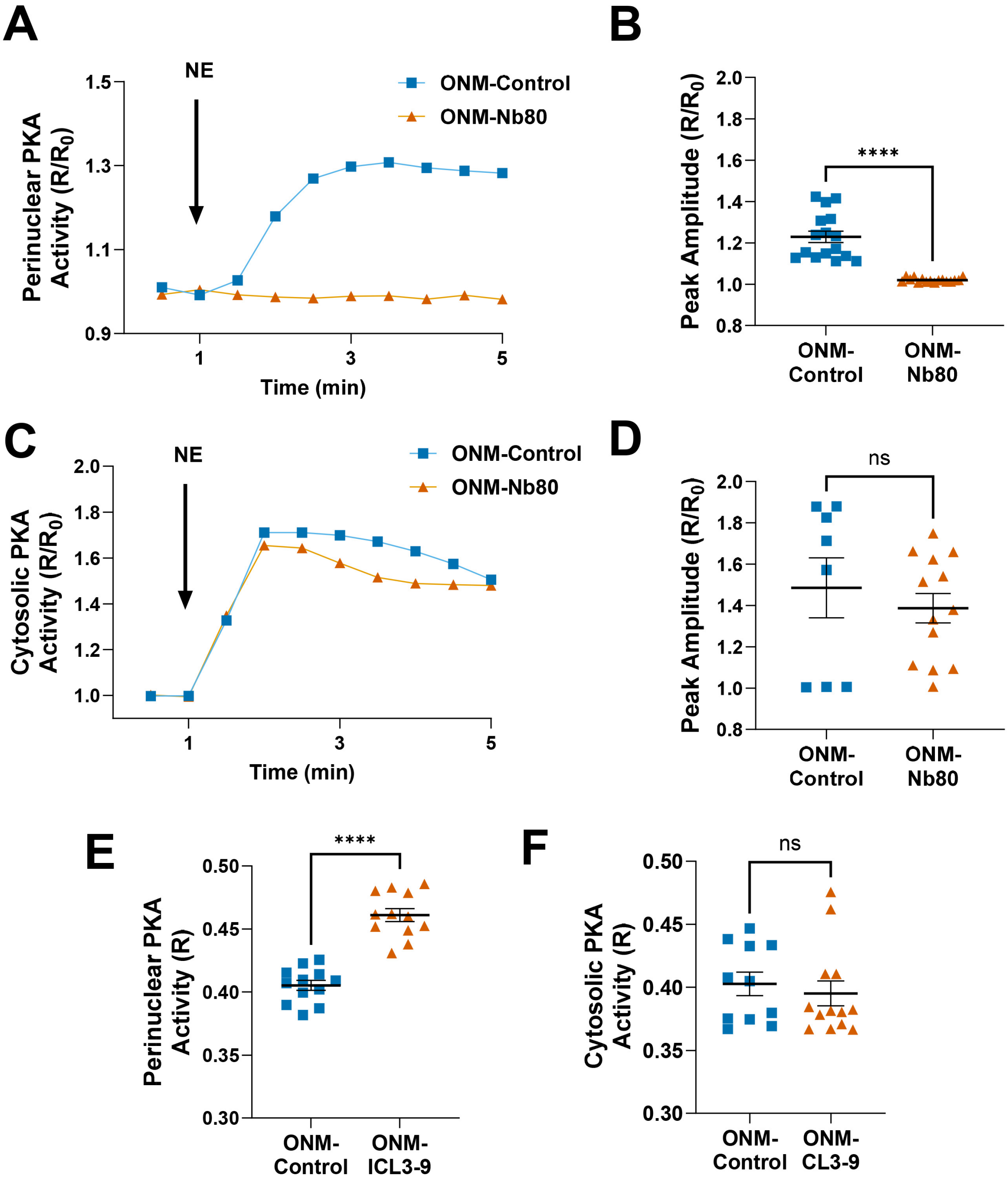

A
NE
ONM-Control
ONM-Nb80
Perinuclear PKA Activity (R/R0)
1.5
1.3
1.1
0.9
1
3
5
Time (min)
B
Peak Amplitude (R/R0)
2.0
1.8
1.6
1.4
1.2
1.0
0.8
****
ONM-Control
ONM-Nb80
C
NE
ONM-Control
ONM-Nb80
Cytosolic PKA Activity (R/R0)
2.0
1.8
1.6
1.4
1.2
1.0
1
3
5
Time (min)
D
Peak Amplitude (R/R0)
ns
ONM-Control
ONM-Nb80
E
Perinuclear PKA Activity (R)
0.50
0.45
0.40
0.35
0.30
****
ONM-Control
ONM-ICL3-9
F
Cytosolic PKA Activity (R)
ns
ONM-Control
ONM-CL3-9

**Supplemental Figure 2: Perinuclear β-adrenergic receptors are necessary and sufficient for activation of AKAP6β-bound PKA in adult rat cardiomyocytes.**

(A-D) FRET imaging of adult rat ventricular myocytes expressing either AKAR4 or ONM-AKAR4 and ONM-Nb80 (top middle) or ONM-Control. Representative tracings and peak amplitudes for FRET ratio (R normalized to baseline $R_0$) are shown.

(E-F) FRET imaging of baseline PKA activity ($R_0$) in adult myocytes expressing either AKAR4 or ONM-AKAR4 and either ONM-Control or ONM-ICL3-9. P-values by T-test.

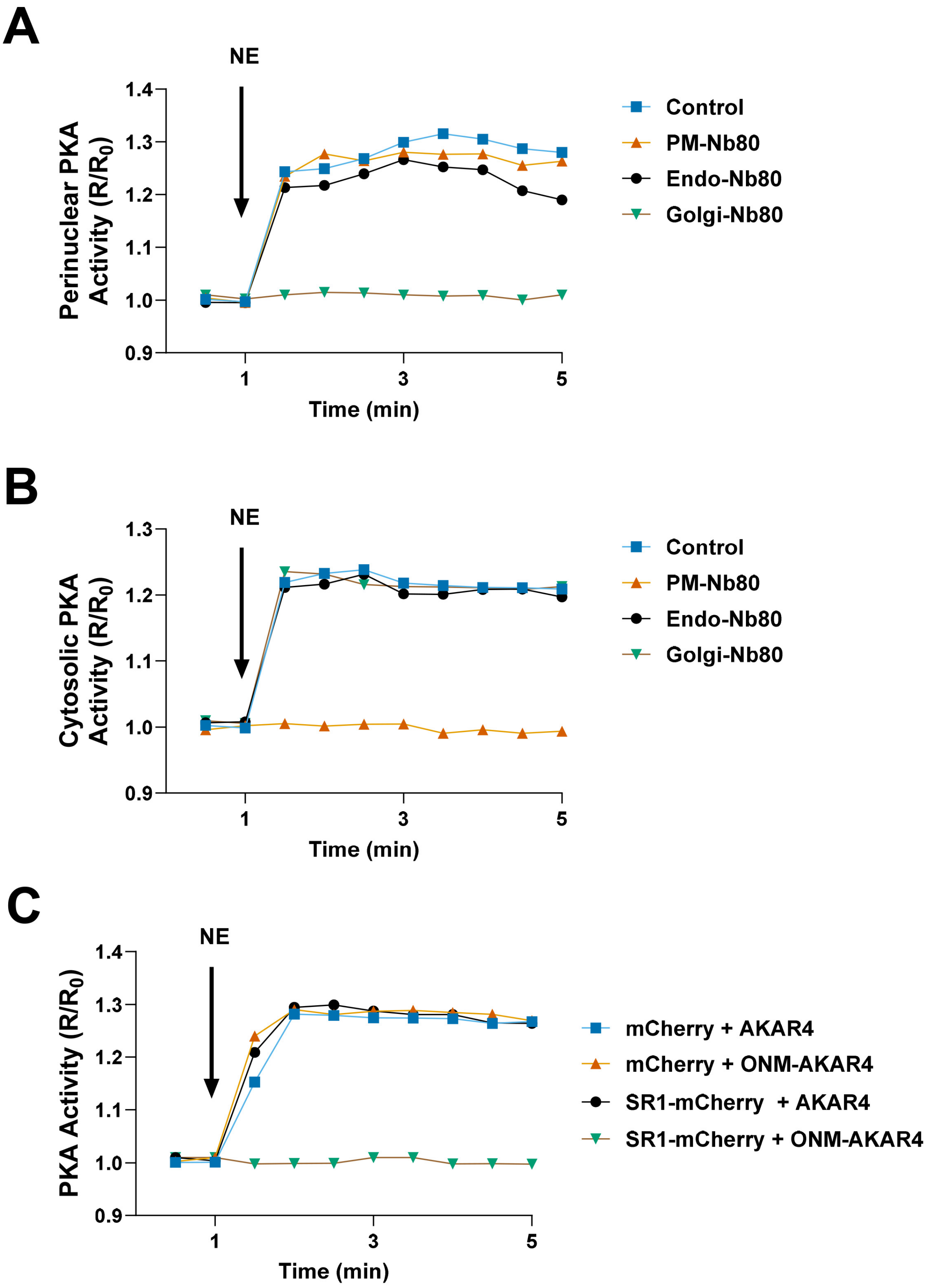
A
NE
Perinuclear PKA Activity (R/R0)
1.4
1.3
1.2
1.1
1.0
0.9
1
3
5
Time (min)
Control
PM-Nb80
Endo-Nb80
Golgi-Nb80
B
NE
Cytosolic PKA Activity (R/R0)
1.3
1.2
1.1
1.0
0.9
1
3
5
Time (min)
Control
PM-Nb80
Endo-Nb80
Golgi-Nb80
C
NE
PKA Activity (R/R0)
1.4
1.3
1.2
1.1
1.0
0.9
1
3
5
Time (min)
mCherry + AKAR4
mCherry + ONM-AKAR4
SR1-mCherry + AKAR4
SR1-mCherry + ONM-AKAR4

**Supplemental Figure 3: Representative tracings for Figure 3.**

(A) See Figure 3C.

(B) See Figure 3D.

(C) See Figure 3H.

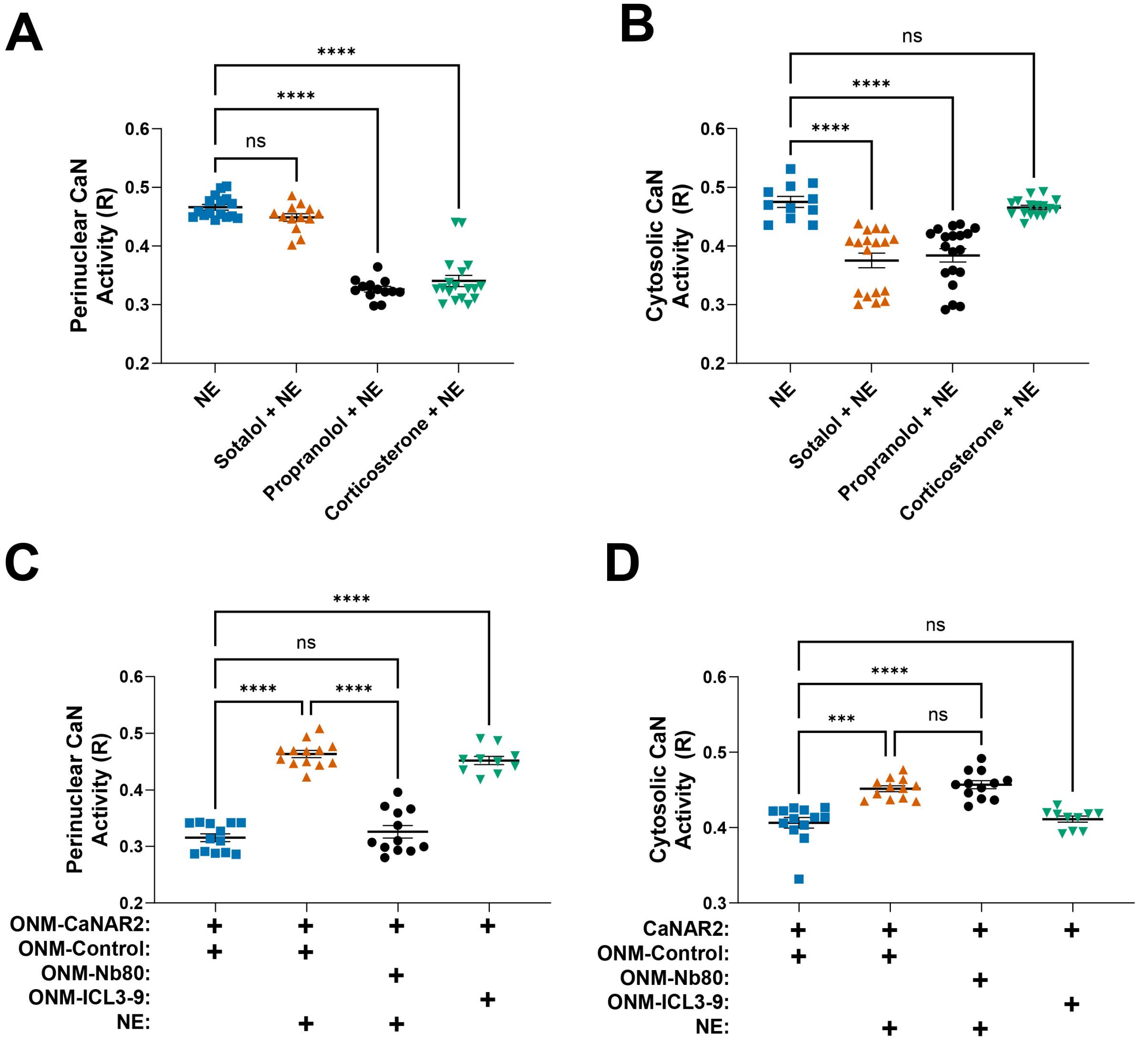


**Supplemental Figure 4: Regulation of $Ca^{2+}$-dependent calcineurin signaling by AKAP6β-associated βARs in adult rat ventricular myocytes.**

CaN activity was measured in adult myocytes expressing CaNAR2, ONM-CaNAR, ONM-Control, ONM-Nb80 and/or ONM-ICL3-9 and treated for 1 day with NE and the indicated inhibitors. CaN activity is reported as FRET ratios (R).

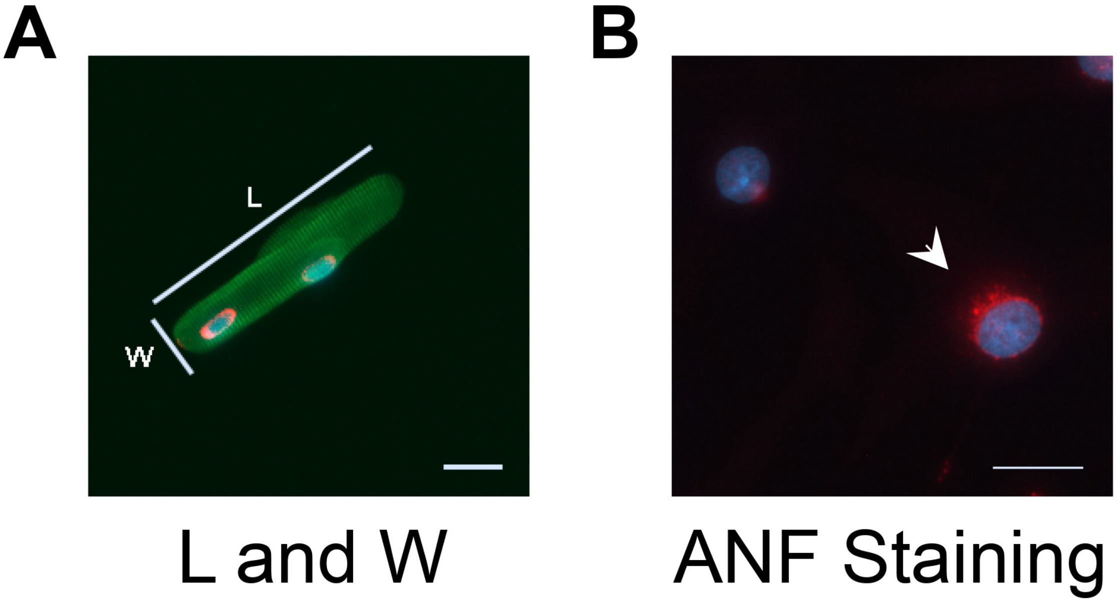


**Supplemental Figure 5: Regulation of myocyte hypertrophy by perinuclear β-adrenergic receptors.**

(A) Example of an adult myocyte expressing ONM-Nb80 (red) stained with α-actinin antibody (green) and Dapi nuclear stain (blue). Length and width measurements are based upon the maximum dimensions perpendicular and parallel to α-actinin sarcomeric staining.

Bars – 20 $\mu$m.

(B) Example of neonatal myocyte with positive ANF staining (red), Dapi nuclear staining in blue.

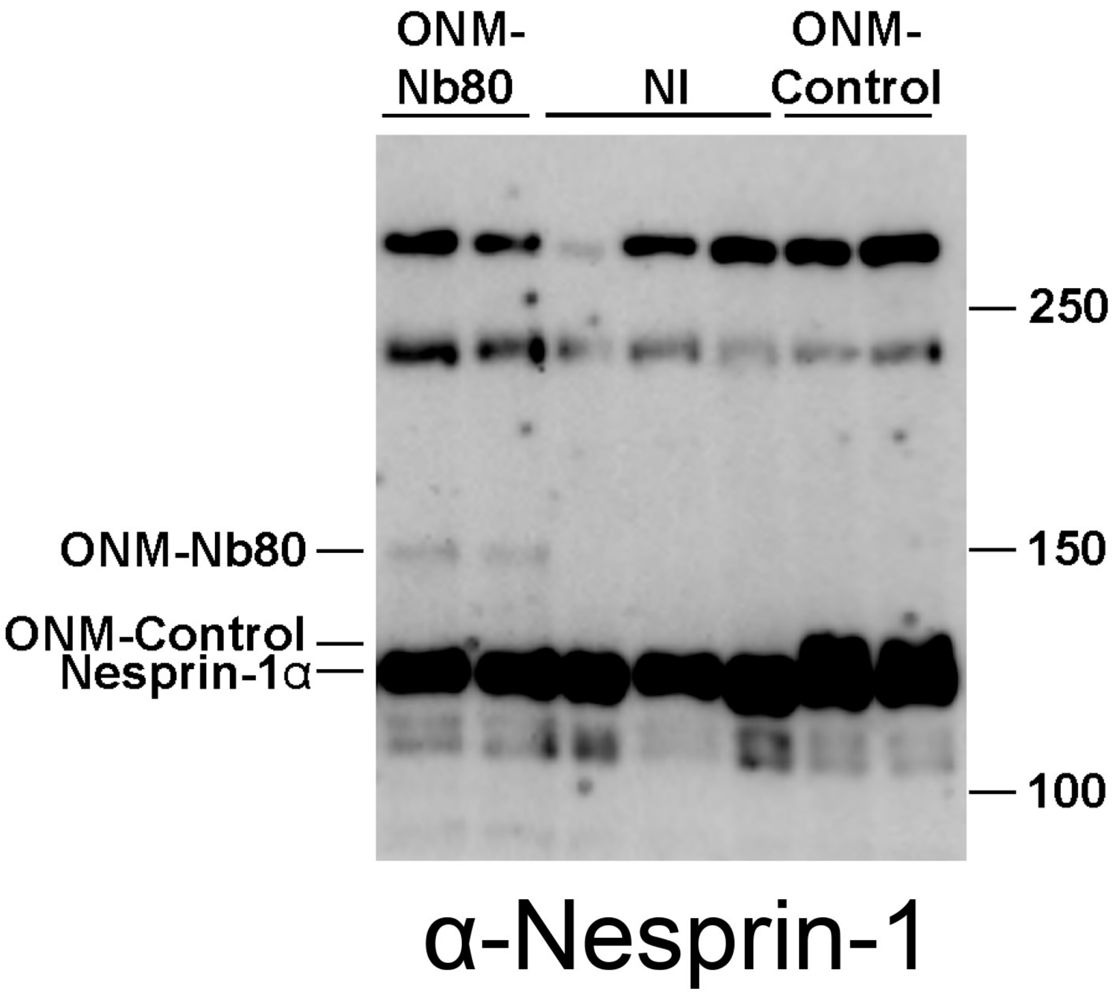


**Supplemental Figure 6: In vivo expression of nesprin-fusion proteins.**

Western blot of NTG mice from TM54 experiment injected developed with α-nesprin-1 antibody. ONM-control migrates slightly slower in SDS-PAGE than the major cardiac isoform nesprin-1α.

1. Rajan, S., Ahmed, R.P., Jagatheesan, G., Petrashevskaya, N., Boivin, G.P., Urboniene, D., Arteaga, G.M., Wolska, B.M., Solaro, R.J., Liggett, S.B., and Wieczorek, D.F. (2007). Dilated cardiomyopathy mutant tropomyosin mice develop cardiac dysfunction with significantly decreased fractional shortening and myofilament calcium sensitivity. Circ Res *101*, 205-214. 10.1161/CIRCRESAHA.107.148379.
2. Cole, N.B., Smith, C.L., Sciaky, N., Terasaki, M., Edidin, M., and Lippincott-Schwartz, J. (1996). Diffusional mobility of Golgi proteins in membranes of living cells. Science *273*, 797-801. 10.1126/science.273.5276.797.
3. Boczek, T., Cameron, E.G., Yu, W., Xia, X., Shah, S.H., Castillo Chabeco, B., Galvao, J., Nahmou, M., Li, J., Thakur, H., et al. (2019). Regulation of Neuronal Survival and Axon Growth by a Perinuclear cAMP Compartment. J Neurosci *39*, 5466-5480. 10.1523/JNEUROSCI.2752-18.2019.
4. Turcotte, M.G., Thakur, H., Kapiloff, M.S., and Dodge-Kafka, K.L. (2022). A perinuclear calcium compartment regulates cardiac myocyte hypertrophy. J Mol Cell Cardiol *172*, 26-40. 10.1016/j.yjmcc.2022.07.007.
5. Nikolaev, V.O., Bunemann, M., Hein, L., Hannawacker, A., and Lohse, M.J. (2004). Novel single chain cAMP sensors for receptor-induced signal propagation. J Biol Chem *279*, 37215-37218. 10.1074/jbc.C400302200.
6. Duong, N.T., Morris, G.E., Lam le, T., Zhang, Q., Sewry, C.A., Shanahan, C.M., and Holt, I. (2014). Nesprins: tissue-specific expression of epsilon and other short isoforms. PLoS One *9*, e94380. 10.1371/journal.pone.0094380.
7. Randles, K.N., Lam le, T., Sewry, C.A., Puckelwartz, M., Furling, D., Wehnert, M., McNally, E.M., and Morris, G.E. (2010). Nesprins, but not sun proteins, switch isoforms at the nuclear envelope during muscle development. Dev Dyn *239*, 998-1009. 10.1002/dvdy.22229.
8. Carr, R., 3rd, Du, Y., Quoyer, J., Panettieri, R.A., Jr., Janz, J.M., Bouvier, M., Kobilka, B.K., and Benovic, J.L. (2014). Development and characterization of pepducins as Gs-biased allosteric agonists. J Biol Chem *289*, 35668-35684. 10.1074/jbc.M114.618819.
9. Ring, A.M., Manglik, A., Kruse, A.C., Enos, M.D., Weis, W.I., Garcia, K.C., and Kobilka, B.K. (2013). Adrenaline-activated structure of beta2-adrenoceptor stabilized by an engineered nanobody. Nature *502*, 575-579. 10.1038/nature12572.
10. Fraser, I.D., Tavalin, S.J., Lester, L.B., Langeberg, L.K., Westphal, A.M., Dean, R.A., Marrion, N.V., and Scott, J.D. (1998). A novel lipid-anchored A-kinase Anchoring Protein

facilitates cAMP-responsive membrane events. EMBO J *17*, 2261-2272. 10.1093/emboj/17.8.2261.

11. Irannejad, R., Pessino, V., Mika, D., Huang, B., Wedegaertner, P.B., Conti, M., and von Zastrow, M. (2017). Functional selectivity of GPCR-directed drug action through location bias. Nat Chem Biol *13*, 799-806. 10.1038/nchembio.2389.
12. Gold, M.G., Lygren, B., Dokurno, P., Hoshi, N., McConnachie, G., Tasken, K., Carlson, C.R., Scott, J.D., and Barford, D. (2006). Molecular basis of AKAP specificity for PKA regulatory subunits. Mol Cell *24*, 383-395. 10.1016/j.molcel.2006.09.006.
13. Prasad, K.M., Xu, Y., Yang, Z., Acton, S.T., and French, B.A. (2011). Robust cardiomyocyte-specific gene expression following systemic injection of AAV: in vivo gene delivery follows a Poisson distribution. Gene Ther *18*, 43-52. 10.1038/gt.2010.105.
14. Pare, G.C., Easlick, J.L., Mislow, J.M., McNally, E.M., and Kapiloff, M.S. (2005). Nesprin-1alpha contributes to the targeting of mAKAP to the cardiac myocyte nuclear envelope. Exp Cell Res *303*, 388-399. 10.1016/j.yexcr.2004.10.009.
15. Li, J., Tan, Y., Passariello, C.L., Martinez, E.C., Kritzer, M.D., Li, X., Li, X., Li, Y., Yu, Q., Ohgi, K., et al. (2020). Signalosome-Regulated Serum Response Factor Phosphorylation Determining Myocyte Growth in Width Versus Length as a Therapeutic Target for Heart Failure. Circulation *142*, 2138-2154. 10.1161/CIRCULATIONAHA.119.044805.
